# Dynamics of Heat Shock Detection and Response in the Intestine of *Caenorhabditis elegans*

**DOI:** 10.1101/794800

**Authors:** Erin K. Dahlstrom, Erel Levine

## Abstract

The heat shock response is the organized molecular response to stressors which disrupt proteostasis, potentially leading to protein misfolding and aggregation. While the regulation of the heat shock response is well-studied in single cells, its coordination at the cell, tissue, and systemic levels of a multicellular organism is poorly understood. To probe the interplay between systemic and cell-autonomous responses, we studied the upregulation of HSP-16.2, a molecular chaperone induced throughout the intestine of *Caenorhabditis elegans* following a heat shock, by taking longitudinal measurements in a microfluidic environment. Based on the dynamics of HSP-16.2 accumulation, we showed that a combination of heat shock temperature and duration define the intensity of stress inflicted on the worm and identified two regimes of low and high intensity stress. Modeling the underlying regulatory dynamics implicated the saturation of heat shock protein mRNA production in defining these two regimes and emphasized the importance of time separation between transcription and translation in establishing these dynamics. By applying a heat shock and measuring the response in separate parts of the animals, we implicated thermosensory neurons in accelerating the response and transducing information within the animal. We discuss possible implications of the systemic and cell level aspects and how they coordinate to facilitate the organismal response.

## Introduction

Proteostasis is a key regulatory process in organisms that maintains the balance between protein synthesis, folding and assembly, and degradation (Balchin *et al*, 2016; Wolff *et al*, 2014). Failure to sustain proteostasis is a hallmark of several important human protein conformational diseases including Alzheimer’s, Parkinson’s, and some types of cancer (Powers *et al*, 2009; Mendillo *et al*, 2012; Morimoto, 2008). In addition to protein misfolding diseases, stressful environmental and internal cues such as high temperatures, metabolic stress (Garcia *et al*, 2007; Raynes *et al*, 2012), and aging (Garigan *et al*, 2002) also disrupt proteostasis, as well as having other negative effects on cellular processes, such as DNA damage (Kantidze *et al*, 2016).

The heat shock response (HSR) is a highly conserved molecular response whose main goal is to prevent protein misfolding and aggregation that can lead to interference with cellular function (Ashburner & Bonner, 1979; Feder & Hofmann, 1999; Morimoto, 2011). After a stressor or disruption, it is activated to restore proteostasis (Ohama *et al*, 2016; Gidalevitz *et al*, 2011). It also has a housekeeping functionality under normal conditions, helping to maintain protein folding under small thermal fluctuations (Balch *et al*, 2008; Somero, 1995). While the HSR pathway was originally thought to be cell-autonomous, more recent work provides evidence for additional layers of regulation and the involvement of multiple signaling pathways (Guisbert *et al*, 2013; Takeuchi *et al*, 2015; van Oosten-Hawle *et al*, 2013). A better understanding of the hierarchy of regulation of the HSR could lead to novel therapies for disease intervention (Bose & Cho, 2016; Balch *et al*, 2008; Westerheide & Morimoto, 2005; Prahlad & Morimoto, 2009).

Dynamical systems properties of the HSR have been well-studied in single cells, including human cell lines, yeast, and bacteria (Starosta *et al*, 2014). In a single cell, the HSR is thought to involve three main sets of players – at least one heat shock transcription factor (HSF), a class of molecular chaperones called heat shock proteins (HSP), and the diverse group of misfolded proteins themselves. Under normal conditions, HSF is located predominantly in the cytoplasm of a cell in a repressor complex with at least one HSP. Some evidence suggests two highly conserved heat shock proteins, HSP-70 (Shi *et al*, 1998) and HSP-90 (Zou *et al*, 1998; Bharadwaj *et al*, 1999), or both, are often a part of this repressor complex. Misfolded proteins caused by a stressor titrate HSP away from HSF, which then undergoes a series of activation steps, including trimerization and relocation to the nucleus. There, it binds to a heat shock element on DNA (Kline & Morimoto, 1997) and induces the transcription of several HSPs. Some of these HSPs act as molecular chaperones, restoring proteostasis by clearing some misfolded proteins and helping others refold (Vabulas *et al*, 2010). Other HSPs that are transcribed are part of the repressor complex and enact a negative feedback loop that represses the activity of HSF. As the misfolded proteins are cleared and refolded, more HSPs are left free to bind back to HSF and repress further transcription (Guo *et al*, 2001). In addition, genes unrelated to the HSR, which are silenced by miRNAs during the heat shock, return to normal activity (Fukuoka *et al*, 2014).

In single cells, the accumulation of misfolded proteins caused by a stressor is thought to be the major cue that starts the HSR by titrating the HSPs away from HSF (Inoue *et al*, 2012; Leach *et al*, 2012). An accumulation of misfolded proteins can also trigger the HSR in multicellular organisms, as the introduction of polyglutamine aggregates in *C. elegans* was shown to induce the HSR without an increase in temperature (Satyal *et al*, 2000). Questions remain, however, as to other ways the response is regulated in multicellular organisms and how externally applied heat leads to the activation of HSF.

In *C. elegans*, previous research implicates the thermosensory AFD neurons and AIY interneurons in the systemic regulation of HSF-1-dependent HSR. The AFD and AIY are necessary for both thermotaxis and thermosensation (Mori & Ohshima, 1995a; Luo *et al*, 2014). They also play a role in regulating the longevity of *C. elegans* at different growth temperatures (Lee & Kenyon, 2009). Mutants with genetic ablations of either neuron were shown to have a HSR that produced 5 - 10 times less HSPs than wild type, following a heat stress (Prahlad *et al*, 2008). The HSR could also be triggered in the absence of heat with the optogenetic stimulation of the AFD neuron, which enhances serotonin release (Tatum *et al*, 2015).

Activation of downstream HSR genes has also been shown to be tissue specific in *C. elegans* (Ma *et al*, 2017; van Oosten-Hawle *et al*, 2013; Guisbert *et al*, 2013). Since the proteome can vary greatly between tissues and cells, the regulatory network of a multicellular organism must be able to allow for different modes of activation within individual cells and tissues (Rieger *et al*, 2006). These cell-non-autonomous levels of regulation are not only shown in relation to the HSR; they have also been seen in the regulation of other stress responses including both the endoplasmic reticulum and mitochondrial unfolded protein responses (Taylor & Dillin, 2013; Durieux *et al*, 2011) and hypoxic and oxidative stress responses (Leiser *et al*, 2016; McCallum *et al*, 2016).

In this paper, we ask how activation of the HSR is coordinated within a tissue of a multicellular organism by studying the activation dynamics of HSP-16.2, a small molecular chaperone expressed throughout the intestine of *C. elegans*. We show that these dynamics stem from a combination of cell-autonomous response and systemic signaling. Quantitative features of the dynamics at different temperatures and durations of heat shock suggest a model in which transcription and translation of heat shock proteins are separated in time. This model attributes the two observed regimes of heat shock response, corresponding to low and high intensities, to the saturation of HSP mRNA production. Model predictions for a temperature-dependent response to repeated heat shocks are tested experimentally. We show that the AFD thermosensory neurons contribute to accelerating the response to heat shock, while the AIY interneurons contribute to the transfer of information across the body, though neither is required for activation of the HSR. We discuss possible implications of the different systemic and cell-autonomous aspects of the HSR and how they coordinate to facilitate the organismal response.

## Results

### Longitudinal measurements of heat shock response dynamics with high time resolution

To characterize the regulatory dynamics of HSR in *C. elegans*, we sought to measure the induction dynamics of downstream heat shock proteins in individual worms under well-defined stress conditions. HSR gene expression dynamics in *C. elegans* are conventionally studied by exposing a population of worms crawling on solid media to a warm environment using either a water bath or an incubator. At each time point, a sub-population of animals is taken off the plates and the level of the gene or genes of interest is assayed, using either biochemical techniques or imaging of anesthetized animals on a microscope slide. These are population-level approaches, precluding the possibility of following the dynamics of activation in individual animals (Link *et al*, 1999; Prahlad *et al*, 2008; Mendenhall *et al*, 2012; Guisbert *et al*, 2013; Mendenhall *et al*, 2015).

To facilitate long-term, high-frequency, longitudinal imaging, namely measurements that preserve the identity of each worm throughout an experiment, we used the previously-described microfluidics device WormSpa (Fig. 1A). Worms were confined to individual chambers and fed with a continuous flow of *E. coli* strain OP50 at a fixed density. Under normal conditions, these worms showed a minimal decrease in size, but were otherwise unstressed and exhibited expected physiological cues including movement, egg laying, and pumping (Fig. EV1A - B, **Vid. EV1**) (Kopito & Levine, 2014). A water channel is located directly above the worm chambers, separated from them by a thin layer of PDMS, allowing for the delivery of precise, temperature-controlled heat shock pulses (HS). In each experiment, synchronized worms were raised to early adulthood, loaded to the device, and given two hours to acclimate before beginning data collection.

**Figure 1.**
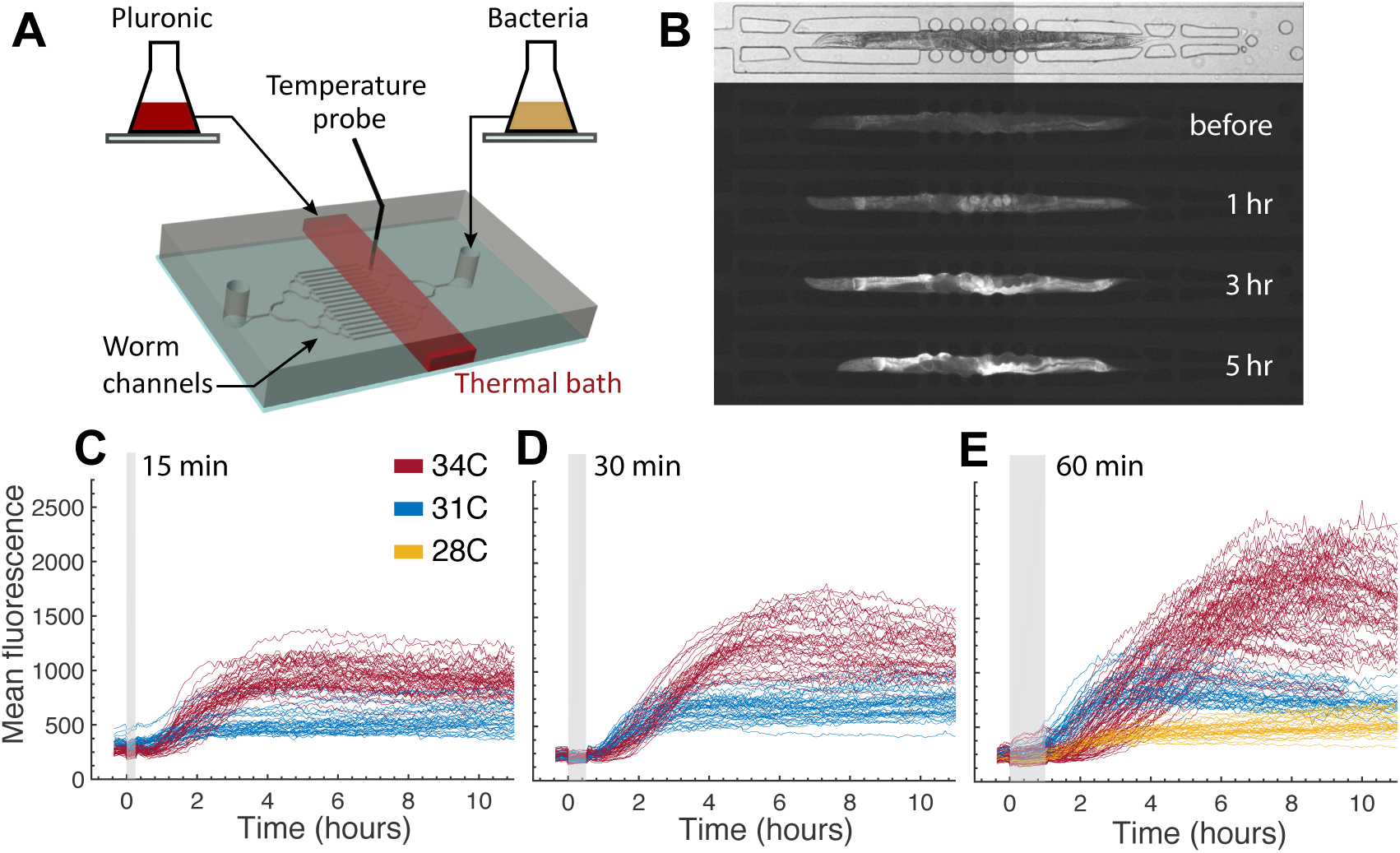
Longitudinal measurements of HSR dynamics with high time resolution. **A** Two-layer microfluidics device. Worms are loaded into bottom channels in bacterial suspension. Heat shock pulse is applied via thermal bath layer located above worms. Temperature is controlled with in-line heater and measured with thermistor probe. **B** GFP fluorescence measured in hsp-16.2p::gfp *C. elegans* maintained in microfluidics devices. Example worm from top to bottom: phase image before heat shock (HS), GFP fluorescence image before 60 minute heat shock pulse at 34°C, GFP 1 hour after HS, GFP 3 hours after HS, GFP 5 hours after HS. **C, D, E** Worms subjected to heat shock pulse in microfluidics device for (**C)** 15 m, **(D)** 30 m, or **(E)** 60 m at 28°C (yellow), 31°C (blue), or 34°C (red). Each line indicates the mean fluorescence of an individual worm. Heat shock pulse is indicated by grey bar. Mean fluorescence was tracked for 10 hours following the heat shock pulse.

The dynamics of HSP-16.2 expression in the intestine were tracked by imaging transgenic worms carrying a transcriptional reporter that drives the expression of GFP from the promoter of a small heat shock protein, hsp-16.2p::gfp, integrated into the worm genome in a single copy (Mendenhall *et al*, 2012). At room temperature (22°C), worms expressed a basal level of GFP fluorescence that then increased on the timescale of hours during and after of a period of elevated temperature (“heat shock pulse”) (Fig. 1B). We applied heat shock pulses of different durations (15, 30, or 60 minutes) and temperatures (28°C, 31°C, or 34°C) and tracked the increase in fluorescence from individual worms for up to 10 hours following the heat shock pulse. Fluorescence measurements in WormSpa following a heat shock pulse are consistent with previously reported measurements performed by the traditional approaches described above (Fig. EV1C - D) (Link *et al*, 1999).

We first considered the dynamics of the total fluorescence signal from each worm under each stress condition (Fig. 1C - E), deferring its distribution across the animal body to later sections. Each curve in this figure corresponds to the mean fluorescence measured from a single animal throughout the experiment. These data suggest that the response dynamics are dependent on both the temperature and the duration of the heat shock pulse.

While the response dynamics we observed were highly consistent and reproducible across worms and experimental repeats, we did observe a clear worm-to-worm variability that was more pronounced at higher temperatures. We hypothesized that this variability may come from higher sensitivity to small changes in temperature across the device. Indeed, at 34°C we observed a significant correlation between the position of a worm in the device and the amplitude of its response, which is not observed at lower temperatures (Fig. EV1E). These data suggest an increased sensitivity to small temperature fluctuations at high temperatures.

### The dynamics of the HSR depend on both temperature and duration of the heat shock pulse

In order to characterize the response dynamics quantitatively, we fit each curve in Fig. 1C - E to a generalized logistic function (see Supporting Text). We used this fit to extract four phenomenological quantities of interest: the time from the start of the heat shock pulse to observed HSR activation (“time lag”); the rate of fluorescence accumulation (“rate”); the fluorescence level at saturation, adjusted to account for the basal level of autofluorescence (“magnitude”); and the rate of decline of GFP fluorescence at the end of an experiment (“decline”) (Fig. 2A). Both the rate of response and magnitude of response increase with increased duration of the heat shock pulse at lower temperatures (Fig. EV1F - G). However, the changes in the rate were more significant at lower temperatures, whereas the changes in magnitude were more significant at higher ones.

**Figure 2.**
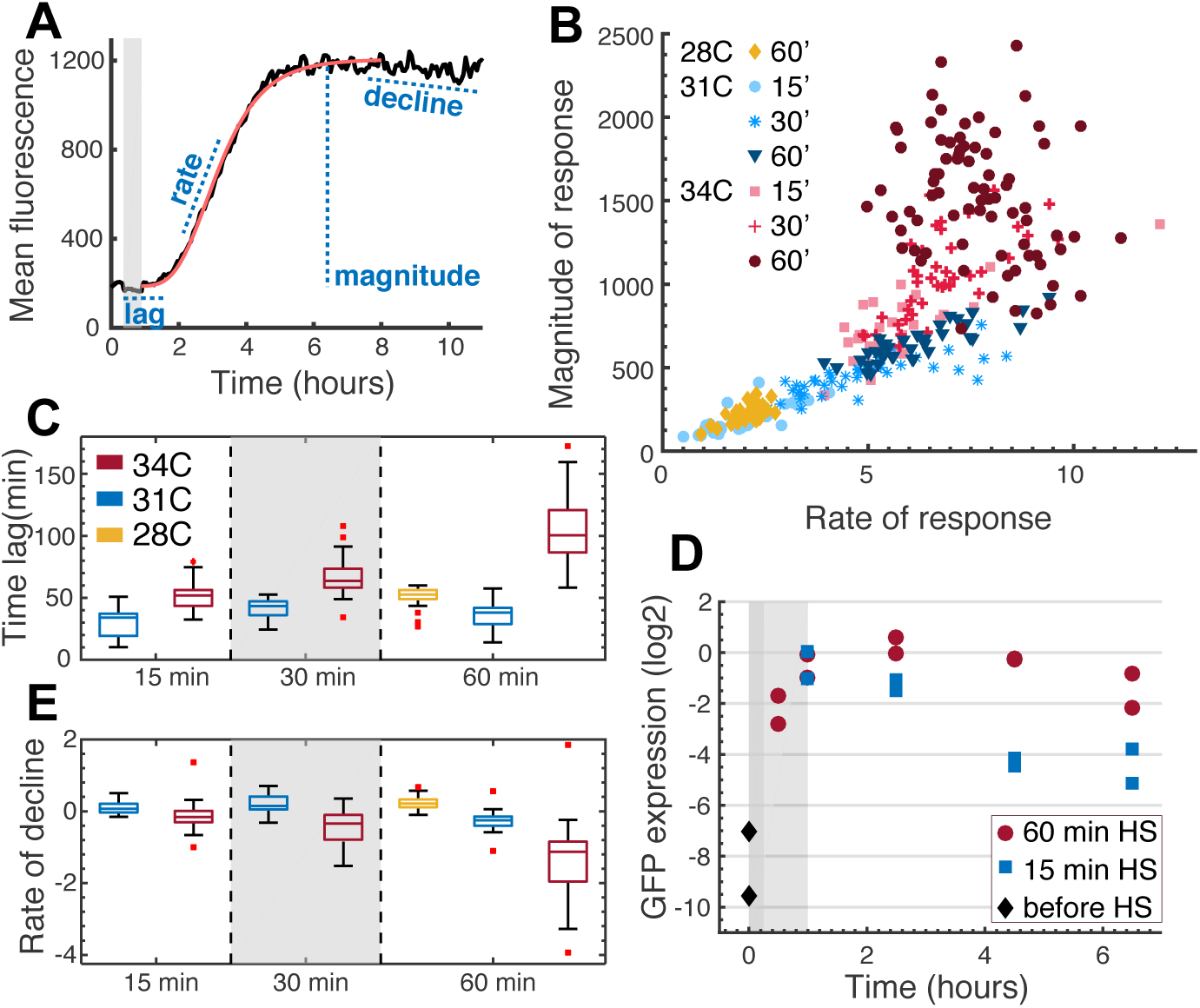
The dynamics of the HSR depend on both temperature and duration of the heat shock pulse. **A** Quantification of HSR dynamics on an example individual worm response curve for a 15 min, 34°C HS. Response is measured as mean fluorescence over whole worm body. Curve is fit to generalized logistic function, then time lag, rate of response, magnitude of response, and rate of decline are analytically calculated. **B** Magnitude of response (mean fluorescence) plotted versus rate of response (mean fluorescence/minutes) for all single heat shock pulses, as indicated in legend. Magnitude is normalized to basal fluorescence for each individual worm at the start of each experiment; rate of response is maximum response rate along logistic fit curve. Points represent individual worms. **C** Time lag of response in minutes, calculated from start of heat shock pulse for all single pulses, as indicated in legend. **D** *log*_2_ GFP mRNA expression for hsp-16.2p::gfp expressing worms as measured by qPCR after 15 minute (blue) or 60 minute (red) heat shock pulse via plates placed in 33°C water bath. Each point is N = 50 worms. Results are normalized to the housekeeping gene *snb-1*. **E** Rate of decline (mean fluorescence/minutes) calculated 8 – 10 hours after heat shock pulse for all single pulses, as indicated in legend.

To explore these differences, we plotted the rate of response against the magnitude of response for individual worms (Fig. 2B). The “data collapse” observed in Fig. 2B, namely the fact that measurements from different animals under different experimental conditions fall on a single curve, suggests that the response dynamics are dictated by a “stress intensity”. This intensity can be described as an integration of both duration and temperature of the heat shock pulse, such that the same intensity leads to the same response. Moreover, this figure suggests that over a range of heat shock pulses (termed “low intensity”), there is a strong linear correlation between the rate and magnitude. Pulses with higher temperature or longer duration (“high intensity”) break from this pattern. Linear correlation between rate at which fluorescence accumulates and the accumulated magnitude suggests that in the low intensity regime, the duration of accumulation is fixed.

Under most stress conditions, we observed that accumulation of fluorescence commenced after the end of the heat shock pulse. This time lag between the environmental shift and the response was temperature-dependent. At lower temperatures (28°C, 31°C), the time lag lasted around 30 - 40 minutes, measured from the start of the heat shock pulse (Fig. 2C), irrespective of the duration of the pulse. In contrast, at 34°C the time lag was longer for a longer heat shock pulse. Interestingly, we observed that at this temperature, the time lag ended 40 minutes after the *end* of the heat shock pulse, irrespective of its duration. (Fig. EV1H).

Previous reports suggest that translation may be inhibited during heat shock (Shalgi *et al*, 2013; Su *et al*, 2016). We therefore asked if the observed lag in HSP-16.2p::GFP accumulation reflects a post-transcriptional block, or a delay in the accumulation of hsp-16.2p::gfp transcripts. To address this question, we quantified the total concentration of hsp-16.2p::gfp mRNA by qPCR (Fig. 2D). Worms were subjected to a 15 or 60 minute heat shock pulse in a 33°C water bath, which qualitatively approximated the fluorescence response from a 34°C heat shock pulse in the microfluidics chip (Fig. EV1C - D). No time lag was observed in the accumulation of mRNA, even for a high intensity stress. Similar results were obtained for the mRNA of hsp-70 (Fig. EV1I), suggesting that the accumulation dynamics of the hsp-16.2p::gfp transcripts are not dominated by its fusion to the GFP-coding region. While we cannot rule out GFP maturation as the cause of the 30- to 40-minute time lag at low temperatures, we consider below the possibility that the additional lag at 34°C lasting the length of the heat shock is due to translational pausing.

GFP is expected to be highly stable in the worm intestine (Dietz & Rief, 2004). This was reflected in the data at low intensities, where we observed little to no decline in the GFP signal up to 10 hours following a heat shock pulse. In contrast, significant decline in the fluorescence signal was observed hours after a pulse classified as high intensity (60 minute HS at 31°C and a 34°C HS of any duration) (Fig. 2E).

### Simple mathematical model provides insight into the mechanisms that drive the dynamics of the HSR

To interpret our results in the context of the regulatory dynamics of the HSR, we developed a mathematical model of the cellular HSR. This model was inspired by previously published models of HSR in mammalian cells (Rieger *et al*, 2005; Scheff *et al*, 2015; Petre *et al*, 2011), but it is considerably simplified, as discussed in the Supporting Text. The model describes the dynamics of three main groups of molecules in the HSR pathway: the heat shock transcription factor (HSF), the heat shock proteins (HSP), and the misfolded or unfolded proteins (Fig. 3A).

**Figure 3.**
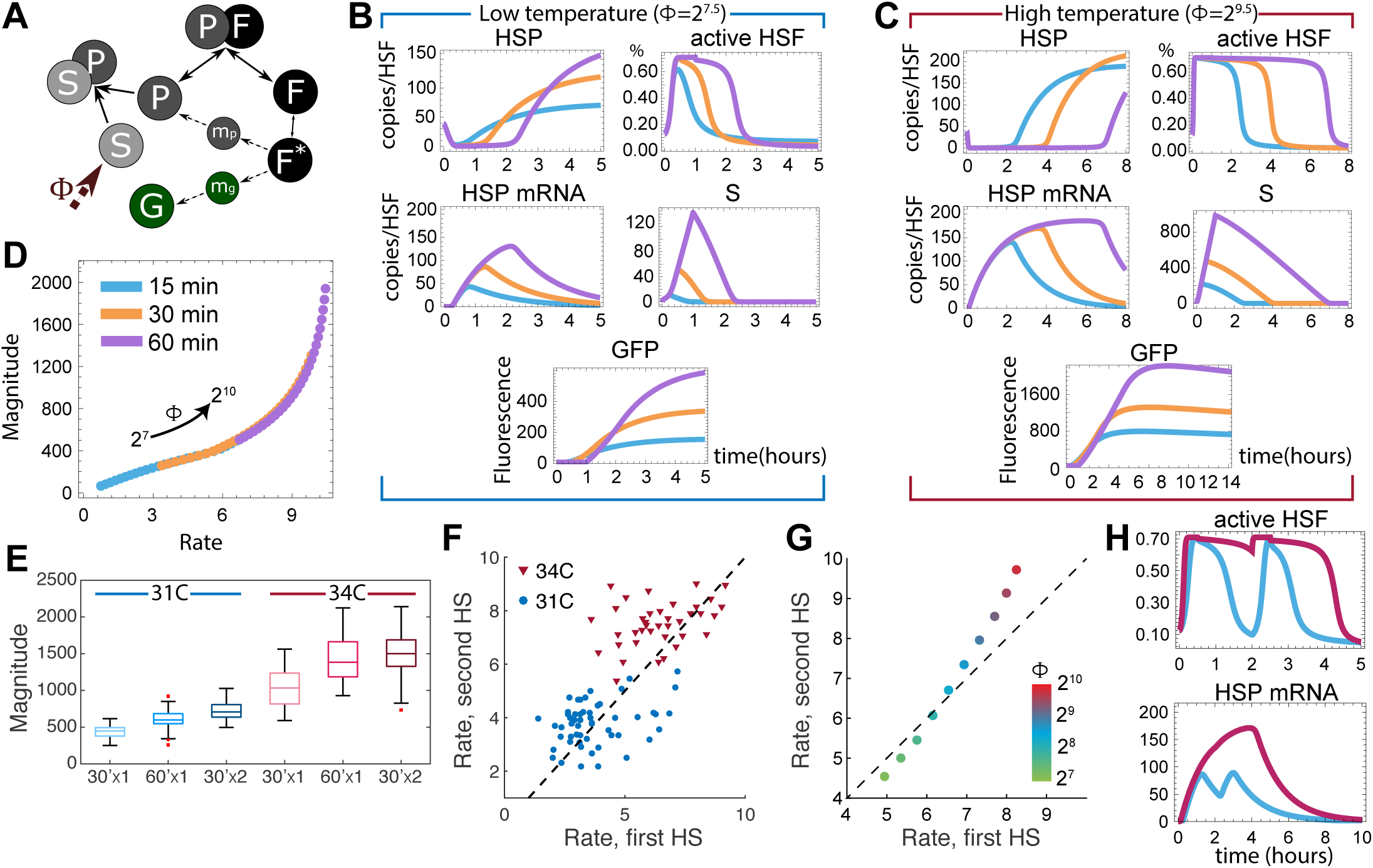
Simple mathematical model provides insight into the mechanisms that drive the dynamics of the HSR. **A** A simple model of the cellular HSR. F is inactive HSF; F* is active HSF; m_p_ and P are mRNA and protein of HSP, respectively; m_g_ and G are the same for GFP; and S is misfolded protein. P and F exist both in a complex and separately. F* generates m_p_ and m_g_, which then generate P and G. A stressor, Φ, generates S, which is sequestered by P. **B, C** Kinetics of P (HSP), p (HSP mRNA), F* (active HSF), S (misfolded proteins), and G (GFP) as predicted by model for a 15 (blue), 30 (orange), or 60 (purple) minute heat shock pulse at **(B)** a low temperature and **(C)** a high temperature. All concentrations are proportional to the total number of HSF, which is set to 1. **D** Magnitude of response (mean fluorescence) plotted versus rate of response (mean fluorescence/minutes) as predicted by model for single heat shock pulses of 15 (blue), 30 (orange), or 60 (purple) minutes over a range of temperatures. **E** Magnitude of response (mean fluorescence) for two consecutive 30 minute heat shock pulses separated by a 90 minute rest (dark color) compared to one single 30 minute (light color) or 60 minute (medium color) heat shock pulse at 31°C (blue) or 34°C (red). **F** Rate of response (mean fluorescence/minutes) for two consecutive 30 minute heat shock pulses separated by a 90 minute rest. Rate of response to second HS plotted versus rate of response to first at 31°C (blue) and 34°C (red). Dotted line indicates first rate of response equal to second rate of response. **G** Rate of response (mean fluorescence/minutes) as predicted by model for two consecutive 30 minute heat shock pulses separated by a 90 minute rest. Pulses are over a range of stress intensity (temperature). Dotted line indicates first rate of response equal to second rate of response. **H** Kinetics of active HSF and HSP mRNA as predicted by model for two consecutive 30 minute heat shock pulses at 31°C (blue) and 34°C (red), separated by a 90 minute rest.

We first assumed that the total cellular concentration of HSF (*F*) in its different forms, *F_total_*, is conserved (Sarge *et al*, 1993). Furthermore, we utilized the fact that the kinetics of binding of HSF to its various partners is rapid compared with other kinetic rates in the system, to write

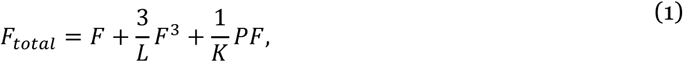

where the first term on the right hand side accounts for the monomeric, inert form of HSF-1; the second to its activated, trimerized form; and the third to HSF-1 in complex with HSP (*P*). In this equation, *L* is the ratio of the inactivation rate of HSF to its activation rate, and *K* is the ratio of the dissociation rate of the HSF:HSP complex to its association rate.

Given the observation of a significant time lag between the accumulation of HSP mRNA and proteins, the model explicitly considers the concentration of both types of molecules, *m_p_* and *P* respectively. We hypothesize that the observed time lag is due to translational inhibition during the heat shock pulse. Under this hypothesis, our observations in Fig. 1C - E suggest that at the lower temperatures, translation resumes towards the end of the heat shock pulse, while at the highest temperature, it only resumes after the pulse. Still, for simplicity we include translational inhibition in our model through a time-dependent rate *λ*(*t*) which takes some value before and after the heat shock and is set to zero during the heat shock. The kinetics of these two molecular species is then given by

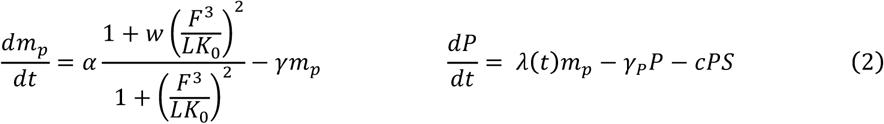

These equations account for the activation of transcription by the active, trimerized form of HSF-1 and assume that the misfolded proteins (*S*) irreversibly titrate HSP proteins, with the association rate of an HSP and a misfolded protein given by *c*. Two similar equations account for the GFP mRNA (*m_g_*) and GFP protein (*G*), with the assumption that our reporter gene is transcribed identically to the HSP itself after a heat shock pulse, but with no basal transcription rate (*α*). In this equation, *K*_0_ is the affinity of activated HSF to the hsp promoter, w is the maximal fold of activation, and *γ* and *γ_P_* are the degradation rates of the mRNA and protein respectively.

Finally, the cellular concentration of misfolded or unfolded proteins is dictated by the rate at which they occur during heat shock, Φ(*T, t*), and by their titration of HSPs, as given by

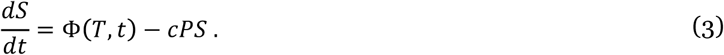

We model Φ(*T, t*) as a step function whose amplitude reflects the temperature of the heat shock pulse and whose width reflects the duration. This is comparable to the heat shock pulses experienced by worms in the microfluidics device, as temperature changes at the start and end of the pulse within 1 - 2 minutes. Lastly, we assume that these misfolded proteins are cleared through a process that both requires and titrates HSPs (the terms proportional to the parameter *c* in Equations 2 and 3). The latter assumption can be weakened without changing our results. A detailed derivation of this model can be found in the Supporting Text.

Definitions of the different variables and parameters of this model and their assigned values can also be found in the tables of the Supporting Text. Some of the parameters were estimated based on either experimental measurements during HSR in *Drosophila* (Fritsch & Wu, 1999; Velazquez *et al*, 1983) and human cell lines (Finka *et al*, 2015; Sarge *et al*, 1993; Wang *et al*, 1999) or our kinetic data. Other parameters simply define the scale of the observable variables and have no impact on our results. The range of amplitudes of the step function Φ (*T, t*) is taken to match the range of observed behaviors, including ratios of GFP saturation between different duration and temperature heat shock pulses and the timescale at which the mean GFP fluorescence saturates.

Our model provides an interpretation of our experimental results. Full activation of the HSR requires exhausting the basal concentration of HSPs that exist in the cell under normal conditions, followed by activation of HSF and accumulation of the HSP transcripts synthesized *de novo*. The time required to exhaust this initial pool of HSPs and the time required to activate all HSFs are both shorter at higher temperatures (Fig. 3B - C). In our model, accumulation of HSPs only starts when the heat shock pulse ends. The concentration of HSP mRNAs at that time sets the rate at which new HSPs are synthesized and accumulated, *i.e.* the measured rate of fluorescence accumulation. The concentration of misfolded proteins at the same time sets the total number of HSPs to be consumed throughout the remaining response, which (due to the stability of GFP) corresponds to the measured fluorescence magnitude.

Thus, when the heat shock pulse is in the low intensity range, both concentrations are proportional to the level of stress at the end of the pulse, which results from a combination of temperature and duration of the pulse. This means that the rate and magnitude of response are linearly proportional to each other (Fig. 3D), as observed experimentally (Fig. 2B). This linearity breaks if the heat shock pulse falls in the high intensity range, where the rate of accumulation of HSP mRNAs is saturated before translation begins. In this case, the rate of HSP accumulation takes its maximal value, and synthesis of the required number of HSPs necessitates a prolonged period of response.

Further confirmation for this mechanism comes from observed long-term decline of the fluorescent signal at high intensities. HSP transcription terminates only when there are enough free HSPs to titrate all HSFs, namely when HSPs are no longer required. Following a strong heat shock pulse, the level of HSP mRNA at this point may still be considerable, leading to further production of HSPs that are no longer needed. This “overshoot” is followed by a noticeable decline despite the fact that the half-life of these proteins may be considerably longer than the duration of the experiment (Fig. 3C). Higher induction of the HSR is thus linked to a higher rate of decline, as observed experimentally (Fig. 2E).

To test the limits of our single negative feedback model, we asked what it predicts about habituation and memory effects. We exposed worms to two 30 minute heat shock pulses, both at either 31°C or 34°C, separated by 90 minutes (Fig. EV2A - D). At both temperatures, the magnitude of the response was additive, resembling that of a single 60 minute pulse (Fig. 3E). At 31°C, the rate of response to the second pulse was similar to that of the first, and both were comparable to the rate of response to a single 30 minute pulse at 31°C. In contrast, at 34°C the rate of response to the second pulse was significantly larger than to the first pulse in most animals, reaching rates comparable to those of a single 60 minute pulse at 34°C (Fig. 3F, Fig. EV2E). These results confirm the predictions of our model, which predicts similar rates in response to two low temperature heat shock pulses and an increase in rate between consecutive pulses at higher temperatures (Fig. 3G). The model can be used to provide an interpretation of these results via the predicted kinetics of different molecules. At lower temperatures, the 30 minute pulse is short enough to avoid saturation of active HSF, and consequently the 90 minute interval between pulses permits complete annihilation of misfolded proteins and a reset of the active HSF to basal level. In contrast, at higher temperatures, HSF remains active in the interval between pulses, continuing production of HSP mRNA and saturating the rate of HSP mRNA production during the second heat shock pulse (Fig. 3H).

### Spatial characteristics of the expression of HSP-16.2 in the worm intestine

So far, we ignored potential differences among the cells of the intestine and considered the level of HSP-16.2p::GFP as its average expression throughout the entire intestine. However, HSP-16.2 is expressed in 20 cells of the worm intestine which are organized into 9 rings that form the intestinal tube. We next sought to characterize possible differences among these cells and how they might depend on the stress intensity. The distribution of the fluorescence signal in two worms, given a 60 minute heat shock pulse at 28°C or 34°C, is depicted in Fig. 4A (and is representative of the dynamics observed in most worms). Every row in this figure comes from a single frame, ordered in time from top to bottom. The intensity of each pixel corresponds to the observed fluorescence at the related position along the long axis of the worm, averaged over the short axis. Lines were aligned by performing spline interpolation and aligning the leftmost and rightmost peaks in intensity. Close inspection of these figures raises the possibility that at 34°C, the farther a cell is from the anterior end, the more delayed the increase in fluorescence is. In contrast, at lower temperatures fluorescence appears to increase simultaneously at all positions along the worm.

**Figure 4.**
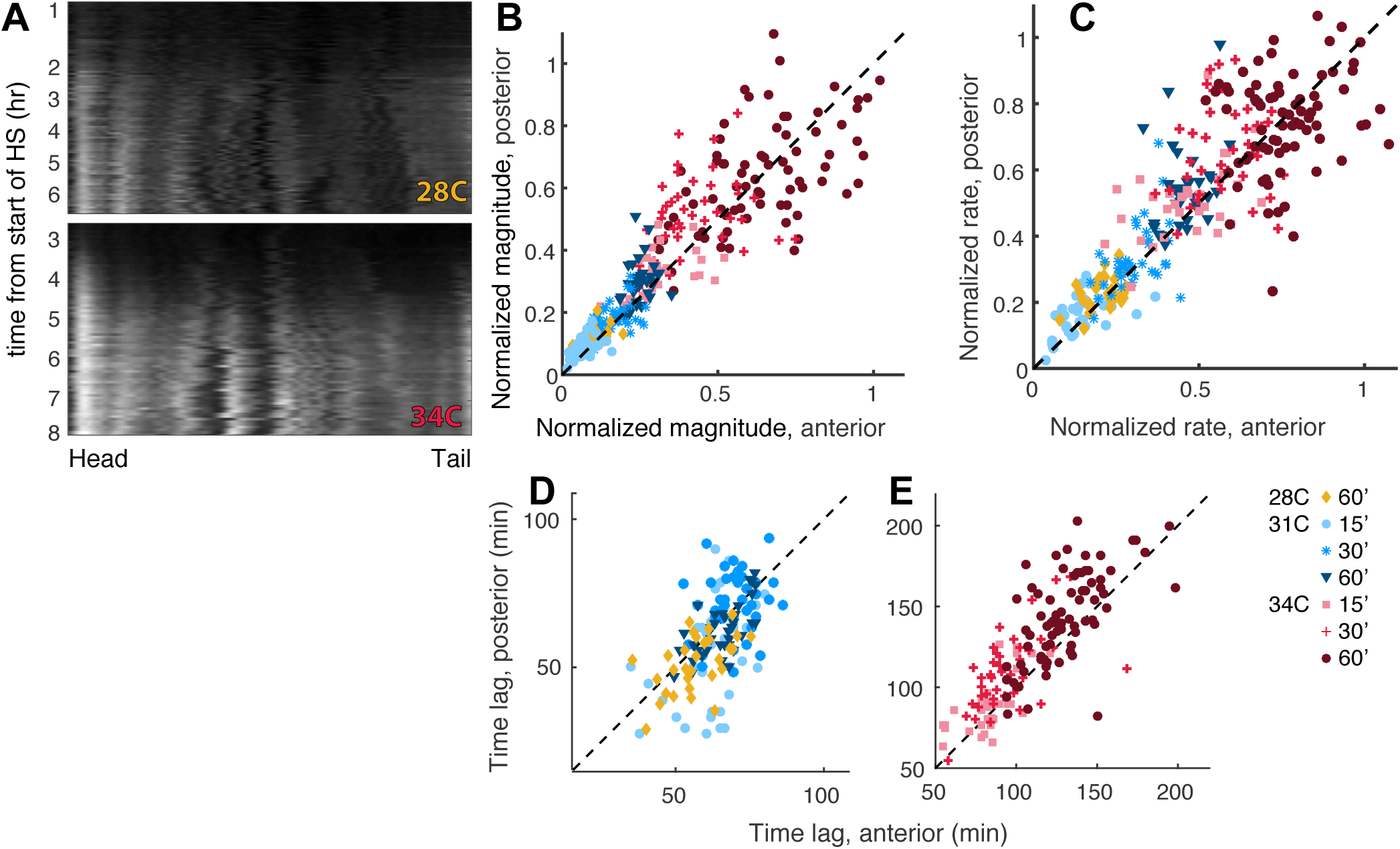
Spatial characteristics of the HSP-16.2 heat shock response. **A** Example time courses for a worm exposed to a 60 minute heat shock pulse at 28°C (top) or 34°C (bottom). Each horizontal row is the mean fluorescence of the worm body at a single time point, collapsed to a single row of pixels. Time is ordered from top to bottom for each time course. **B, C (B)** Magnitude of response (mean fluorescence) and **(C)** rate of response (mean fluorescence/minutes) of posterior part of worms plotted versus anterior part of worms subject to a whole-body, single heat shock pulse, duration and temperature as indicated in legend. Dashed black line is y = x. Magnitude and rate are both normalized to their respective maximum mean values in the anterior and posterior thirds. **D, E** Time lag (minutes) from start of heat shock pulse for posterior part of worms plotted versus anterior part of worms subject to a whole-body, single heat shock pulse at a **(D)** low temperature or **(E)** high temperature; duration and temperature are as indicated in legend. Dashed black line is y = x.

To explore this observation and quantify it, we partitioned the intestine into three equal parts: anterior, middle, and posterior. The middle section of the intestine overlaps with the uterus, and in some cases, embryos *in utero* express a fluorescent signal which interferes with the signal from the intestine of the mother. We therefore ignored this third and focused on the differences between the anterior and posterior cells. To do this, we plotted the mean GFP fluorescence in each part separately (Fig. EV3) and parameterized these curves using a logistic fit, as above.

We then asked if the dependence of the magnitude and rate of response on the stress intensity is the same in both parts (Fig. EV4A - D). To factor out differences in the measured fluorescence that come from the variable numbers and sizes of cells in each third, we normalized the measured fluorescence for each worm’s anterior and posterior by the maximum observed mean fluorescence in the anterior or posterior. After normalization, we found that both the magnitude and rate of response in the two sections are highly correlated (R = 0.88 and 0.84, respectively, Fig. 4B - C). This was also the case for the time lag of the activation for a heat shock pulse at 28°C or 31°C (Fig. 4D). In contrast, the time lag after a 34°C pulse was significantly longer in the posterior for all pulse durations (Fig. 4E, Fig. EV4E). This suggests that the observed propagation of the fluorescence signal from head to tail (Fig. 4A) is due to a delay in the initiation of response and not a modulation of its dynamics once it starts.

### Thermosensory neurons may accelerate intestinal response to heat shock

The observation that the HSR starts earlier in the anterior of the intestine than in the posterior raises the hypothesis that some signal propagates from the head to the tail. One possible source of such a signal is a pair of thermosensory neurons located in the head. Two likely candidates are the AFD thermosensory neurons and AIY interneurons, both of which have been previously implicated in thermotactic behavior (Mori & Ohshima, 1995a) and linked to the HSR (Prahlad *et al*, 2008). To investigate the potential roles of these neurons, we characterized the HSR dynamics in mutant worms carrying either the *ttx-1 (p767)* allele, which prevents terminal differentiation of the AFD neurons (Satterlee *et al*, 2001), or the *ttx-3 (ks5)* allele, which ablates the functionality of the AIY interneurons (Hobert *et al*, 1997). The dynamics of HSP-16.2p::GFP accumulation in individual mutant worms under a variety of stress conditions are shown in Fig. EV5.

Neither the activation of the HSR nor the linear relationship between the magnitude and rate of response at low intensity stresses requires either the AFD or AIY neurons (Fig. 5A - B). However, while the break in linearity at high intensity stresses in the AIY-ablated mutants was similar to that in wild type animals, in the AFD-ablated animals, it occurred at lower stress intensities. At the higher intensities, these worms respond with a lower rate than wild type but reach a higher magnitude. These observations also hold separately for the anterior and posterior thirds of the intestine in both mutants (Fig. EV6A - D).

**Figure 5.**
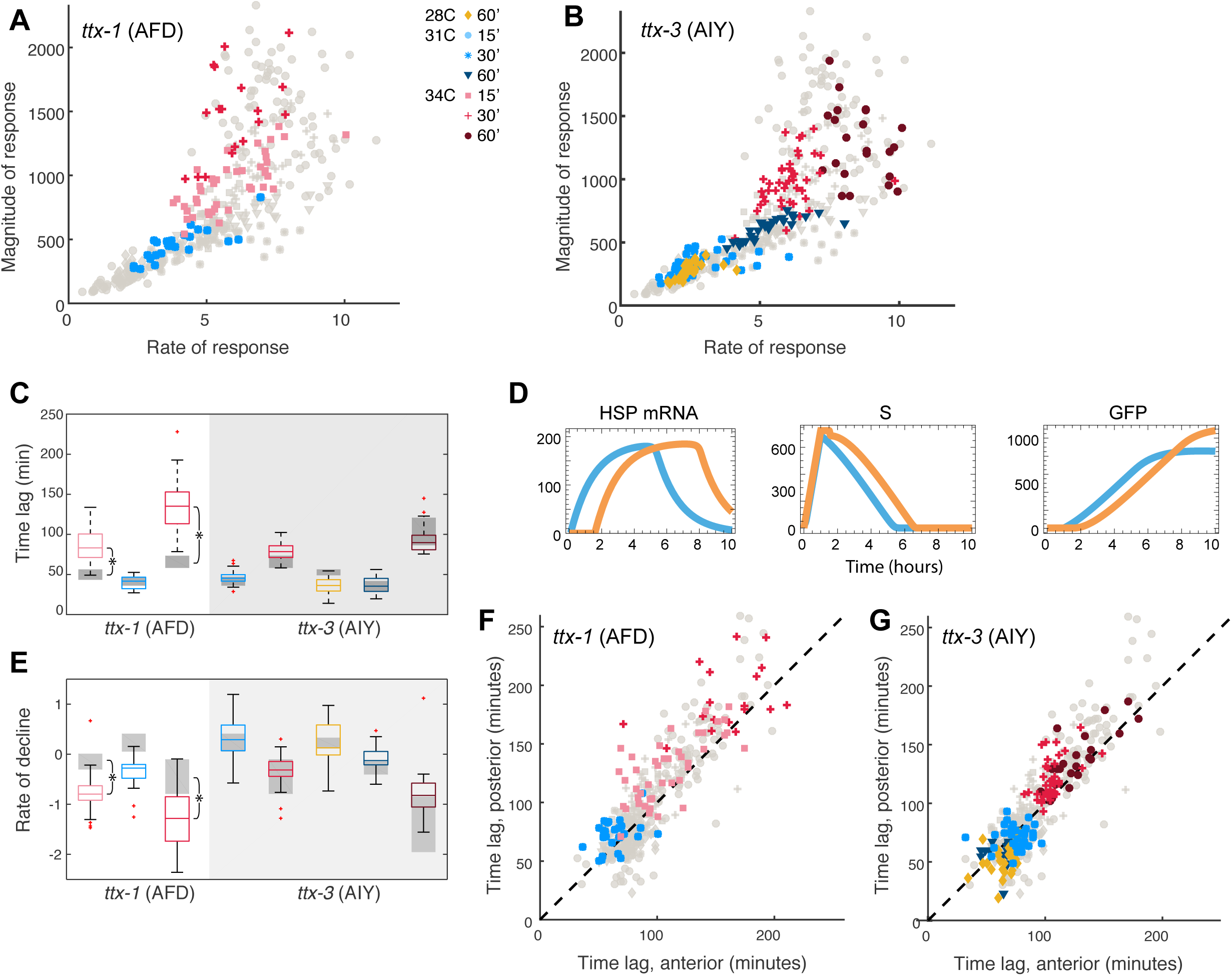
Thermosensory neurons set a threshold for maximum rate of response of the HSR. **A, B** Magnitude of response (mean fluorescence) plotted versus rate of response (mean fluorescence/minutes) for all single heat shock pulses, duration and temperature as indicated in legend, for **(A)** *ttx-1* (AFD) mutants and **(B)** *ttx-3* (AIY) mutants. Magnitude is normalized to basal autofluorescence of each individual worm at start of each experiment; rate of response is maximum response rate along logistic fit curve. Points represent individual worms. Light grey points are wild type worms of the corresponding experimental condition. **C** Time lag of response in minutes, calculated from start of heat shock pulse for all single heat shock pulses for *ttx-1* (AFD) and *ttx-3* (AIY) mutants. Temperature and duration are color-coded, as indicated by the legend. Grey boxes represent the corresponding experiment in wild type worms. Statistically significant differences between mutant and wild type worms are indicated by an asterisk: for 15 and 30 minute HS at 34°C, p-values are <10^-11^ and <10^-14^, respectively. **D** Kinetics of HSP mRNA, misfolded proteins (S), and GFP as predicted by model for a single 60 minute heat shock pulse at 34°C (blue) and the same heat shock pulse with a 30 minute delay (orange). **E** Rate of decline (mean fluorescence/minutes) calculated 8 – 10 hours after heat shock pulse ends for all single heat shock pulses, duration and temperature as indicated in legend, for *ttx-1* (AFD) and *ttx-3* (AIY) mutants. Grey boxes represent wild type worm data. Statistically significant differences between mutant and wild type worms are indicated by an asterisk: for 15 and 30 minute HS at 34°C, p-values are <10^-3^ and <10^-10^, respectively. **F, G** Time lag (minutes) from start of heat shock pulse of posterior part versus anterior part of **(F)** *ttx-1* (AFD) and **(G)** *ttx-3* (AIY) mutant worms subject to a whole-body, single heat shock pulse; duration and temperature as indicated in legend.

In search of possible mechanisms behind the change in dynamics in AFD-ablated animals, we turned to our model and tested the hypothesis that AFD-dependent signaling alters one or more of the kinetic rates of this model. However, the relationship between magnitude and rate, as depicted in Fig. 3D, turned out to be robust to changes in parameters within the relevant range (estimations in the Supporting Text). This comes from the fact that parameter changes lead to proportional effects on both the level of HSP mRNA, *m_p_*, (which in turn sets the rate) and on the demand for HSPs by *S* (which sets the magnitude). Thus, if the two were linearly proportional with the original set of parameters, they remained proportional after the parameter change.

Looking for an alternative explanation for the modified relationship between magnitude and rate, we considered the time lag before activation. At low stress intensities, the time lag of both the *ttx-1* and *ttx-3* mutants was similar to that of the wild type animals. At high intensities, however, the time lag was significantly prolonged in AFD-ablated mutants but not in AIY-ablated mutants (Fig. 5C). For example, the time lag for a 30 minute HS at 34°C in an AFD-ablated worm surpassed that of a 60 minute HS in a wild type worm. Motivated by this observation, we used our model to test the hypothesis that a delayed start of the response may lead to the observed increase in magnitude with no increase in the rate. To do this, we introduced a time delay between the emergence of misfolded proteins and their detection by HSPs (see Supporting Text for details). A direct consequence of this change to the model is a prolonged time lag before activation. This leads to a delay in the kinetics of HSF and HSP during heat shock but does not change the accumulation of active HSFs or the level of HSP mRNA (Fig. 5D). The observed rate of HSP accumulation is therefore unaltered. In contrast, as the accumulation of misfolded proteins is unchanged by HSP sequestration for a longer period of time, the demand for HSPs, which sets the magnitude of the response, is increased (Fig. 5D). Thus, this model predicts that a delay in the time of detection results in an increased magnitude of response with no effect on its rate, as observed experimentally.

As discussed above, our model links high stress intensities with the observed late decline in fluorescence. Given the overshoot in the response in the AFD-ablated mutants, the model predicts an increase in the rate of decline in these worms as compared with wild type animals. This prediction is confirmed by the experimental data that show a higher rate of decline for the AFD-ablated mutants but not for the AIY-ablated mutants (Fig. 5E).

Lastly, we asked if the thermosensory neurons are involved in the observed propagation of HSR activation from head to tail (Fig. 4A). The dynamics of response, i.e. rate and magnitude, to heat shock in the anterior and posterior thirds were highly similar, as seen in wildtype animals (Fig. EV6E - F). Moreover, our data did not implicate these neurons in generating the time lag difference between the anterior and posterior parts, which still exists in both mutant strains (Fig. 5F - G).

### Non-local activation of the heat shock response is independent of thermosensory neurons

Finally, we asked how activation of the HSR in one part of the intestine depends on the stress experienced by another part of the intestine. To address this question, we modified our microfluidics device to have two water channels, each perpendicular to the worm chambers and positioned above one half of the worm body (Fig. 6A). The temperatures of the two channels were controlled independently, allowing us to apply a 30 minute heat shock pulse at either 31°C or 34°C to one half of the worm body while maintaining the other half at room temperature. We then tracked the dynamics of HSP-16.2p::GFP accumulation in both halves (Fig. EV7). Surprisingly, we find an increase in fluorescence in both halves of the worm, including the half that was maintained at room temperature.

**Figure 6.**
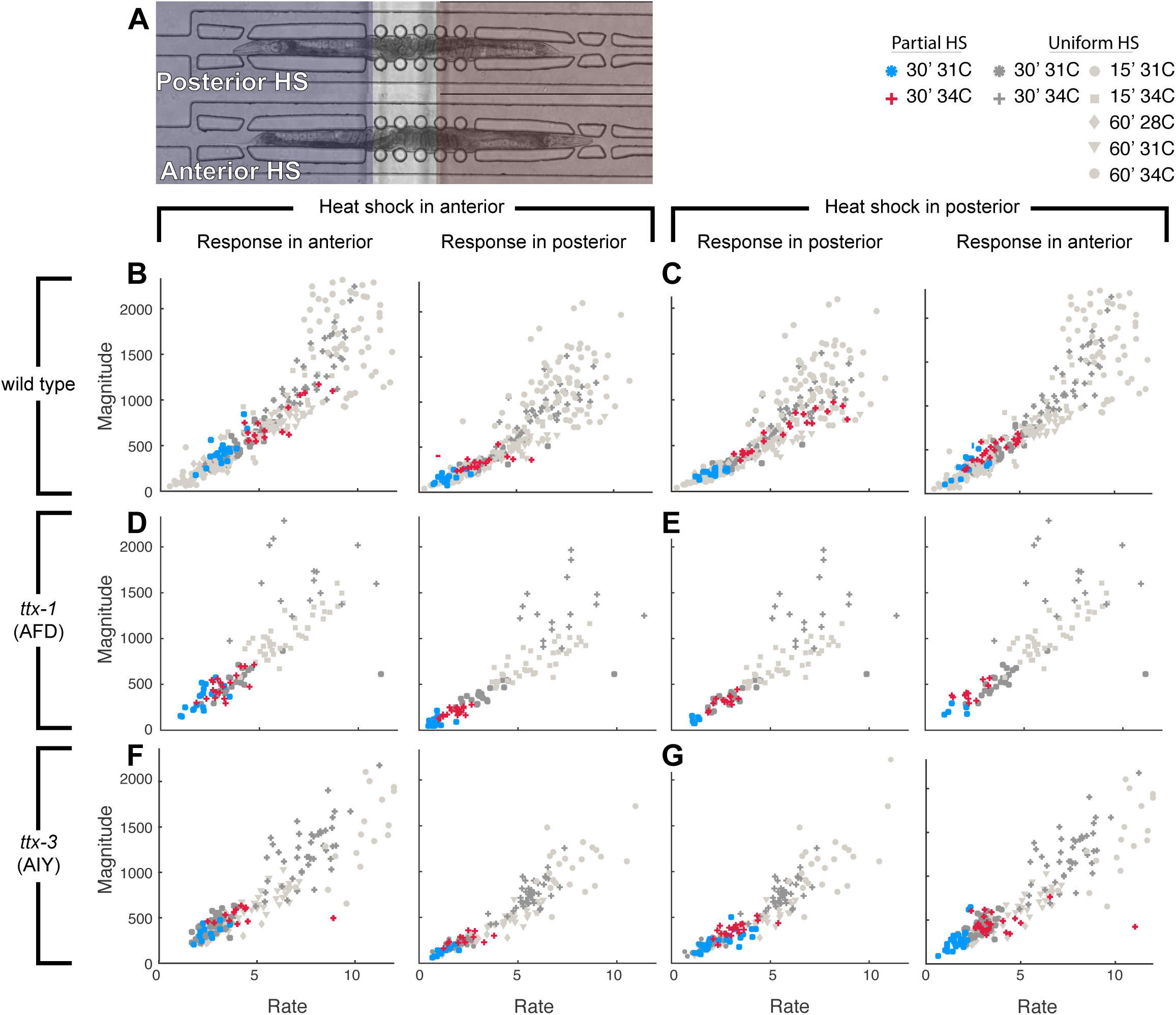
Linearity of rate and magnitude is independent of thermosensory neurons. **A** Variation on microfluidics device from Figure 1A with an additional water channel separated from the first so their temperatures can be maintained separately. Water channels are positioned over the body of the worm such that one part of the worm – either the anterior or the posterior – can be heat shocked (red) at a time, while the rest of the worm is maintained at room temperature (blue). **B – G** Magnitude of response (mean fluorescence) plotted versus rate of response (mean fluorescence/minutes) for **(B,C)** wild type worms, **(D,E)** *ttx-1* (AFD) mutants, and **(F,G)** *ttx-3* (AIY) mutants for single 30 minute heat shock pulse at 31°C (blue) or 34°C (red). Four columns from left to right are: (1) Heat shock anterior, measure anterior; (2) Heat shock anterior, measure posterior; (3) Heat shock posterior, measure posterior; (4) Heat shock posterior, measure anterior. Points represent individual worms. Dark grey points are relevant wild type, *ttx-1*, or *ttx-3* data points for anterior or posterior third of a whole-body heat shocked worm. Light grey points are data points from other non-relevant wild type, *ttx-1*, or *ttx-3* experiments, heat shock pulse and duration as indicated in legend.

When wild type animals were heat shocked in the anterior, the response in the anterior was substantially suppressed as compared with its response to a similar heat shock pulse to the entire body of the worm. The response in the posterior, while even smaller, was significant (Fig. 6B). In both cases, the relationship between magnitude and rate of response was found within the linear regime of the same “universal” curve observed above. Heat shock applied to the posterior half of the worm resulted in a corresponding pattern: a suppressed response in the posterior and an even more suppressed but still significant response in the anterior (Fig. 6C). These results suggest that induction of the HSR in a cell results not only from the stress on its own proteome, but also from heat-triggered signaling from other cells.

In the context of our mathematical model, these observations suggest that some of the parameters that define the kinetics of HSR are affected by external signaling that carries information about the stress level in other parts of the body. These parameters could be, for example, the basal transcription rate *α* or the transcriptional activation level *w*. The available data, however, does not allow us to distinguish between these possibilities.

To test the involvement of the thermosensory neurons in this cell-non-autonomous response, we repeated these experiments with the two mutant strains from above. When either the AFD or AIY neurons were ablated, the response to a partial heat shock was highly suppressed in both the heat shocked and room temperature parts of the animals (Fig. 6D - G). For the AFD ablation, this behavior was similar to what was observed for a whole-animal heat-shock. In contrast, AIY ablation had a noticeable effect only when worms were partially heat-shocked. This suggests that the AIY neurons may play a role in communicating heat shock information between different parts of the worm body.

Above, we observed that for a 34°C heat shock pulse, the time lag to activation of the response is longer in the posterior than in the anterior and is prolonged in AFD-ablated worms. Both of these features were not observed when the heat shock pulse was applied to only part of the worm (Fig. EV8). The time lag observed following either a 31°C or 34°C heat shock pulse was always around 30 - 40 minutes in both the anterior and the posterior, irrespective of temperature, the part of the body that was heat shocked, or the ablation of the thermosensory neurons.

## Discussion

The heat shock response is a universal program tasked with maintaining the stability and integrity of the proteome under normal and stress conditions. The accepted paradigm posits that regulation of the heat shock response is done predominantly by the heat shock factor(s) through upregulation of the transcription of heat shock proteins in response to conditions that cause protein misfolding. Modulation of the activity of the heat shock factor comes in part from its association with some of the larger heat shock proteins, constituting a negative feedback, but can also come from post-translational modifications that promote nuclear retention and DNA binding (Sarge *et al*, 1993) or DNA dissociation and inactivation (Kline & Morimoto, 1997; Ohama *et al*, 2016).

Over the years, the heat shock response has received much attention from the systems and quantitative biology communities. Many properties make this system particularly attractive for modeling. First, its function is of major biomedical importance, with implications to the survival, fitness, and evolution of organisms from all kingdoms of life. Dysfunction of the heat shock response has consequences in health and disease, as well as implications in oncogenic processes. Second, the heat shock response is an ancient and universal mechanism that is required for the marginal stability of much of the proteome and the exposure of cells to a broad range of temperatures and other proteotoxic environmental and physiological conditions. Third, despite differences in the identity and origins of some of the proteins involved, the core design of the regulatory network that controls the heat shock response, which is based on titration of a master regulator by heat shock proteins, is conserved from bacteria to humans. Finally, this regulatory network is well defined, well separated from other regulatory functions in the cell, and composed of a relatively small number of protein families, making it amenable for modeling. Correspondingly, the heat shock response has been modeled in bacteria (Kang, 2012), yeast (Castells-Roca *et al*, 2011), and HeLA cells (Rieger *et al*, 2005).

Previous models of the heat shock response at the cellular level aimed to characterize the importance and functionality of different aspects of the control circuit, including transcriptional, post-transcriptional and post-translational regulation, as well as the feedforward and feedback loops (Guisbert *et al*, 2008; Castells-Roca *et al*, 2011; Rieger *et al*, 2005; Petre *et al*, 2011; El-Samad *et al*, 2005; Abravaya *et al*, 1991; Krakowiak *et al*, 2017; Kurata *et al*, 2006). These models vary significantly in the level of detail at which different processes are modeled. In bacteria and yeast, the resolution of available experimental data allowed for assignment of functional roles to different branches of the regulatory circuits (El-Samad *et al*, 2005; Kurata *et al*, 2006) and the association of HSP-70 with HSF-1 (Krakowiak *et al*, 2017), respectively. In contrast, models of the heat shock response in HeLA cells rely on lower resolution experimental data (Rieger *et al*, 2005). While some of these models explicitly account for multiple processes in great detail (Abravaya *et al*, 1991), it has been argued that a minimal model that limits the number of degrees of freedom and uses a coarse-grained description of the underlying molecular processes suffices to explain these data (Sivéry *et al*, 2016). In this paper, we demonstrated that a favorable combination of high-resolution experimental data and a minimal model can help in uncovering the essential determinants of the HSR dynamics.

Beyond technical benefits, such as reduced computational effort and lower risk of spurious parameter fitting, minimal models have a clear advantage in that they are more readily interpretable. In addition, complex models with many fitting parameters may be capable of reproducing the observed data even if their underlying assumptions are wrong. Under such conditions, the failure of a minimal model is more evident, and it may also be simpler to identify the incorrect assumption and then formulate and test alternative hypotheses. In the model we developed here, failing to separate the dynamics of HSP transcription and translation resulted in a clear mismatch between model predictions and the observed dynamics.

Previous models that place HSF-1 titration by HSPs at the center of HSR regulation identified three operating regimes of HSR and related them with normal, acute, and chronic stress conditions (Sivéry *et al*, 2016; Sriram *et al*, 2012). These three regimes are characterized by inactivation, partial activation, or saturation of HSP promoters by HSF-1, respectively. The acute and chronic conditions are related to the low and high intensity stresses we observed experimentally and in our model, but our classification relies on global dynamical properties of the response, rather than on instantaneous properties. Importantly, we find that the classification of a heat shock pulse to one of these regimes depends both on its temperature and on its duration.

Interestingly, the binding efficiency of HSPs to HSF (or its functional homologs) was identified as the key determinant of the sensitivity and robustness of the system in gram-positive and gram-negative bacteria (Inoue *et al*, 2012), in the fungus *Candida albicans* (Leach *et al*, 2012), and in HeLa cells (Rieger *et al*, 2005). These results are puzzling in the context of a multicellular organism: while the level of non-sequestered HSPs can be very different in different cells, the affinity of HSPs to HSF-1 cannot be independently tuned in each cell type. Indeed, our model suggests that this parameter has little effect on the dynamics of HSR given the observed fact that at homeostasis, the concentration of HSPs is high enough to titrate all HSF-1 (Wang *et al*, 1999).

Previous studies compared mathematical modeling with experiments that exposed *C. albicans* (Leach *et al*, 2012) or mammalian cells (Cates *et al*, 2011) to two consecutive heat shock pulses. Both studies found that the response to the second pulse depended on the properties (temperature and duration) of the first. We performed two-pulse experiments mainly as a tool for validating our mathematical model, which was constructed based on our single-pulse data. Our experimental data depicts a similar behavior in the intestine of *C. elegans*, and our model suggests that the dependence on the intensity of the stress comes from the fact that the coupling between the two heat shock pulses comes mainly from the degree at which misfolded proteins of the first pulse are cleared before the initiation of the second.

Heat shock is known to evoke transcriptional pausing of non-heat-shock genes. HSPs, in contrast, are actively transcribed, and their mRNAs are preferentially processed and bypass some stages of quality control for quicker export from the nucleus (Zander *et al*, 2016; Niskanen *et al*, 2015; Maxwell *et al*, 2014). In addition, translation is known to be inhibited during heat shock as well through elongation pausing (Bouche *et al*, 1979; Lindquist, 1980; Spriggs *et al*, 2010; Shalgi *et al*, 2013). Consistent with these findings, our data confirms quick transcription of HSPs under high stress intensities but suggests a significant delay in translation of GFP from these mRNAs. Interestingly, we find that at lower temperatures, these proteins accumulate already towards the end of the heat shock pulse, while at higher temperatures translation only significantly resumes after the return to normal temperatures.

The use of microfluidics was instrumental for this study. The WormSpa device permits the administration of precise heat shock pulses, as well as longitudinal observation of individual animals before, during, and after the perturbation. By following individual animals, we were able to relate the rate and magnitude of the response, which led us to observe the universal behavior that defines low intensity stress and the deviation from this behavior at higher intensities, which drove the development of our model. Longitudinal imaging also allowed us to observe a progression of the fluorescence signal along the intestine from the anterior to the posterior. Microfluidics also allowed application of stress to only part of the animal body. Since worms go either head-first or tail-first into the device at roughly equal frequencies, we were able to measure response to anterior- and posterior-only heat shock in the same experiment.

Previously it has been suggested that the AFD thermosensory neurons and the AIY interneurons, which are required for thermotaxis (Mori & Ohshima, 1995b; Luo *et al*, 2014), are essential for activation of heat shock response (Prahlad *et al*, 2008). Here we find that neither of these neurons is required for heat shock response when heat shock is applied to the entire worm body. However, in the absence of functional AFD neurons, the response to high intensity stress was delayed, particularly in the posterior part of the animal. The AFD neurons have ciliated dendrites that are exposed to the external environment; it is possible that sensing temperature directly from the environment accelerates the response to temperature changes in the intestine.

When heat shock was applied to only part of the animal body, response was observed throughout the animal. This response, however, was significantly reduced as compared with the response to a homogeneous pulse of the same temperature, even in cells that were exposed. This suggests that some properties of the cellular HSR depend on external signals that integrate information from multiple loci in the body. Indeed, genetic perturbation to either the AFD or AIY neurons has devastating effects on the response to partial heat shock, implicating the nervous system in regulating such signals. Identifying these putative signals is an important question for a future study. In this context, it may be relevant to note that several signaling systems have been shown recently to be involved in regulating activity of HSF-1 and its downstream targets, including insulin-like signaling (Chiang *et al*, 2012), integrin signaling (Kumsta *et al*, 2014), and serotonin (Tatum *et al*, 2015).

The observation that cells at the anterior end respond to stress earlier than cells at the posterior end can be interpreted in several ways. It is unlikely that this result is an experimental artifact: the heat-carrying fluid flows in a direction transverse to the worm body, and (as mentioned) worms can be situated in the device with their head-tail axis either along or opposite the flow of food, giving no noticeable effect on the HSR dynamics. It is possible that cells respond autonomously to the change in temperature with time lags that increase along the intestine, although it would be unclear what causes such position-dependent time lag. Alternatively, it is possible that a small temperature gradient develops in the intestine from the flow of ingested liquid. Another possible explanation is that some intercellular molecular signal propagates in the same direction and contributes to these activation dynamics. This could be the same signal mentioned in the context of the thermosensory neurons. An interesting possibility for cell-to-cell communication is the transmission of proteins between cells (Nussbaum-Krammer & Morimoto, 2014; Zullo *et al*, 2015). Movement of HSPs between cells has been detected in *Drosophila* and could play a role in *C. elegans* as well (Takeuchi *et al*, 2015).

## Materials and Methods

### Strains

All strains were maintained on standard nematode growth medium (NGM) plates seeded with *E. coli* strain OP50 at 22°C (Stiernagle, 2006). Early adult hermaphrodites were used in all assays. The N2 Bristol strain was used as wild type, in addition to the following strains: TJ3001 [*zSi3001* [*hsp-16.2p::GFP::unc-54 + Cbr-unc-119(+)*] *II*]], ERL35 [*ttx-1 (p767) V; zSi3001 II*], ERL36 [*ttx-3 (ks5) X; zSi3001 II*], PR767 [*ttx-1 (p767) V*],and FK134 [*ttx-3 (ks5) X*].

### Device fabrication

The microfluidics devices used were extended two-layer versions of the previously published WormSpa (Kopito & Levine, 2014). The bottom layer was a 50 µm thick WormSpa layer made with spin coated SU8 3050, baked at 65°C for 2 minutes and 95°C for 20 minutes, then irradiated (exposed to UV) at around 200 mJ/cm^2^ for 30 seconds. Post-exposure, the wafer was baked for 1 minute at 65°C and 4 minutes at 95°C, then developed in PGMEA. The top layer is a 300 µm water channel used for temperature control during the heat shock pulse. After spin coating with SU8 2150, the wafer was soft baked for 8 minutes and 75 minutes at 65°C and 95°C respectively, then UV exposed for 45 seconds. Post exposure bake was 5 minutes and 25 minutes respectively at 65°C and 95°C. Both masks were coated with heptadecafluoro – 1, 1, 2, 2 – tetra-hydrodecyl trichlorosilane by soaking for 10 minutes in a 0.1% solution with Novec 7100 (HFE).

The water channel was made with PDMS mixed in a 10:1 ratio of Sylgard 184 silicone elastomer base to silicone elastomer curing agent (Dow Corning) and separated from the worm channels by a 100 µm membrane. The membrane was created by spin coating the worm channel wafer with two very thin layers of PDMS – the first was 10:1 PDMS spin coated at 150 rpm for 5 seconds followed by 300 rpm for 60 seconds, and the second was 5:1 PDMS spin coated at 375 rpm for 5 seconds followed by 750 rpm for 60 seconds. The worm channel wafer was baked for 30 – 60 minutes at 65°C to partially cure the PDMS, then the water channel was plasma bonded on top. After allowing the whole device to cure for 2 – 12 hours at 65°C, it was carefully removed from the wafer and plasma sealed to a glass slide.

### Microfluidics device experiments

For all microfluidics experiments, an overnight culture of *E. coli* OP50-1 grown in LB media was centrifuged and re-suspended in S-medium (Stiernagle, 2006) to an optical density of 5 at 600 nm (OD_600_ = 5). The bacteria was then heat killed for 35 minutes in a 65°C water bath, as previously described (Gruber *et al*, 2007; Soukup *et al*, 2012) and cooled to room temperature. Heat killed bacteria created less build-up in the microfluidics device than live bacteria, which improved the fluid flow. This method of heat killing was confirmed by plating bacteria and verifying no colony growth. Optical density was measured prior to heat treatment to ensure consistency between experiments. The bacteria were then filtered with a 0.5 µm syringe filter to remove any large particulates that could clog the microfluidics channels.

Age synchronized worms were obtained by letting 20 – 40 gravid adults lay eggs on a seeded plate for an hour, then removing the adult worms. Plates were incubated at 22°C for 60 – 70 hours. 32 – 35 worms were picked from the plate, placed into the prepared bacterial suspension in a 600 µL, non-stick tube and loaded into the microfluidics device as previously described (Kopito & Levine, 2014). Worms were given two hours post-loading to adjust to the device before data acquisition began.

During the experiments, bacterial suspension was delivered to the worm chambers at a rate of 5 µL/min, with 10 second pulses at 200 µL/min applied every 20 minutes to clear bacterial buildup and eggs away from the worms. Flow was controlled with an in-lab developed custom LabView script and the New-Era NE-501 OEM syringe pump. Bacterial solution was contained in 10 mL male luer lock syringes equipped with 20 gauge, ½” length, blunt tip, stainless steel industrial dispensing tips from CML Supply (Item 901-20-050). Medical grade polyethylene microtubing with an inner diameter of 0.86 mm and an outer diameter of 1.32 mm (BB31695-PE/5, Scientific Commodities, Inc.) was used to connect the microfluidics device to the syringes. The syringes and syringe pump were kept on a VWR Standard Analog Shaker at speed setting 5 throughout the experiment to help prevent settling of the bacteria to the bottom of the syringe and ensure fixed density throughout the experiment.

The water channel was connected to a reservoir of 2.5% pluronic solution with the same medical grade polyethylene microtubing. The pluronic solution was filtered with a Corning 500 mL 0.22 µm filter system and driven by gravity flow through the device. A pluronic solution was used rather than water alone to help prevent the collection of bubbles in the water channel which disrupt imaging. A heat shock pulse was applied using the SF-28 in-line heater, placed directly over the tubing right before entry into the microfluidics device, and the TC-324C temperature controller, both from Warner Instruments. Temperature in the water channel was measured with a Physitemp Instruments IT-24P insulated, Type T, Copper-Constantan thermocouple with a polyurethane insulted wire connected to the EXTECH Instruments Process PID Controller 48VFL. The tip of the thermocouple was enclosed in a combination of a syringe tip, a 200 µL pipette tip, and a small piece of BB31695-PE/1 tubing in order to be inserted into the microfluidics device and stay watertight.

### Data acquisition and analysis

Images were acquired using a Zeiss Axio Observer Z1 inverted microscope with a 10X objective and a Hamamatsu Orca II camera with 10 ms phase exposure and 50 ms GFP exposure from a Colibri LED light source. For each condition, we imaged 30 – 83 worms in 2 or more independent repeats. Imaging took place for 20 minutes before the start of a heat shock, during the heat shock, and up to 10 hours post exposure. Each worm was imaged every 2 minutes during the heat shock pulse and every 3 to 5 minutes before and after.

To analyze images, the worm body was identified in each frame and a mask created using morphological image analysis techniques including erosion and dilation. After applying the mask, mean fluorescence over the entire worm body or a fraction of the worm body was calculated, generating an individual worm response curve. Each response curve was then fit to a generalized logistic function and relevant quantities were extracted (see Supporting Text). All image analysis was performed with in-lab developed, custom MATLAB scripts.

### qPCR experiments

Worms for qPCR experiments were synchronized by bleaching, as described (Stiernagle, 2006), plated 12 – 16 hours after the bleaching, and allowed to grow for 55 – 60 hours at 22°C before starting the heat shock. Around 300 worms were grown on each 10 cm plate.

To heat shock the worms, the plates were sealed with parafilm and placed in a 33°C water bath. Plates were weighted down with empty 250 mL glass bottles to ensure they were fully submerged for the length of the heat shock. The plates were then given 2 minutes to equilibrate to the temperature of the water bath, heat shocked for 15 or 60 minutes, and then moved immediately to a room temperature water bath to cool for 5 minutes. The first time point after the heat shock for a 60 minute heat shock was taken immediately after removal from the room temperature water bath.

At each time point, 50 worms were picked into TRI-reagent. RNA was extracted and treated with DNase I (NEB) to remove all DNA from sample. cDNA was synthesized using the ProtoScript® First Strand cDNA Synthesis Kit (NEB). Lastly, qPCR was performed using the KAPA SYBR FAST qPCR Kit. Results were normalized to the housekeeping gene *snb-1*.

## Acknowledgements

We thank Eunju (Lucy) Lee for technical assistance and Neil Peterman and Kyung Suk Lee for discussion. This work was performed in part at the Center for Nanoscale Systems (CNS), a member of the National Nanotechnology Infrastructure Network (NNIN), which is supported by the National Science Foundation under NSF award no. ECS-0335765. Some strains used in this work were provided by the CGC, which is funded by NIH Office of Research Infrastructure Programs (P40 OD010440). EKD was supported by an NDSEG fellowship. This work was supported in part by the National Science Foundation under award no. MCB-1413134.

## Author contributions

EKD and EL designed the research, analyzed the data, developed the model, and wrote the manuscript; EKD performed all experiments.

## Expanded View Figure Captions

**EV Figure 1.**
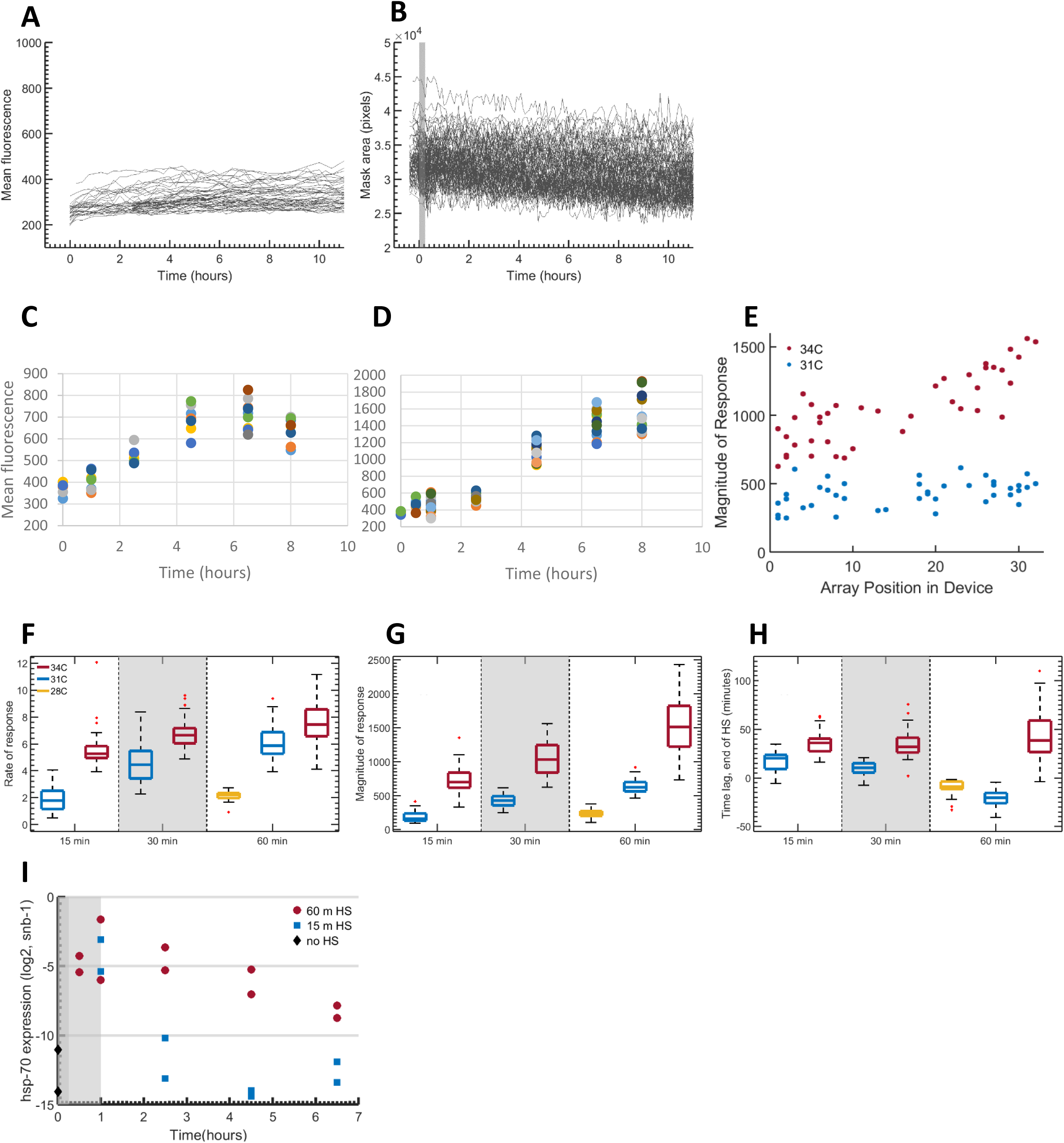
Extended data for single heat shock pulse experiments with wild type worms. **A** Example mean GFP fluorescence response curves for individual worms in microfluidics device with no heat shock pulse applied. **B** Example mask size (pixels) from image analysis pipeline as measure of worm body size throughout experiment for worms exposed to a 15 minute heat shock pulse at 31°C or 34°C. **C, D** Example mean GFP fluorescence response curves for individual worms on plates exposed to a **(C)** 15 minute or **(D)** 60 minute heat shock pulse in a 33°C water bath, as quantified by the traditional method of immobilization of subpopulations of worms on an agar pad at each time point. **E** Magnitude of response for a 30 minute heat shock pulse at 31°C (blue; *R*^2^ = 0.29, p-value = 0.07) or 34°C (red; *R*^2^ = 0.81, p-value < 10^-9^) plotted against position of worm in microfluidics device, where 1 is farthest from in-line heater, and 32 is closest to in-line heater. **F** Rate of response (mean fluorescence/minutes) for all single heat shock pulses, as indicated in legend. Rate is maximum response rate along logistic fit curve. **F** Magnitude of response (mean fluorescence) for all single heat shock pulses, as indicated in legend. Magnitude is normalized to basal autofluorescence of each worm at the start of each experiment. **H** Time lag of response in minutes, calculated from *end* of heat shock pulse for all single pulses, as indicated in legend. **I** *log*_2_ hsp-70 mRNA expression as measured by qPCR after 15 minute (blue) or 60 minute (red) heat shock via plates placed in 33°C water bath. Each point is N = 50 worms. Results are normalized to the housekeeping gene *snb-1*.

**EV Figure 2.**
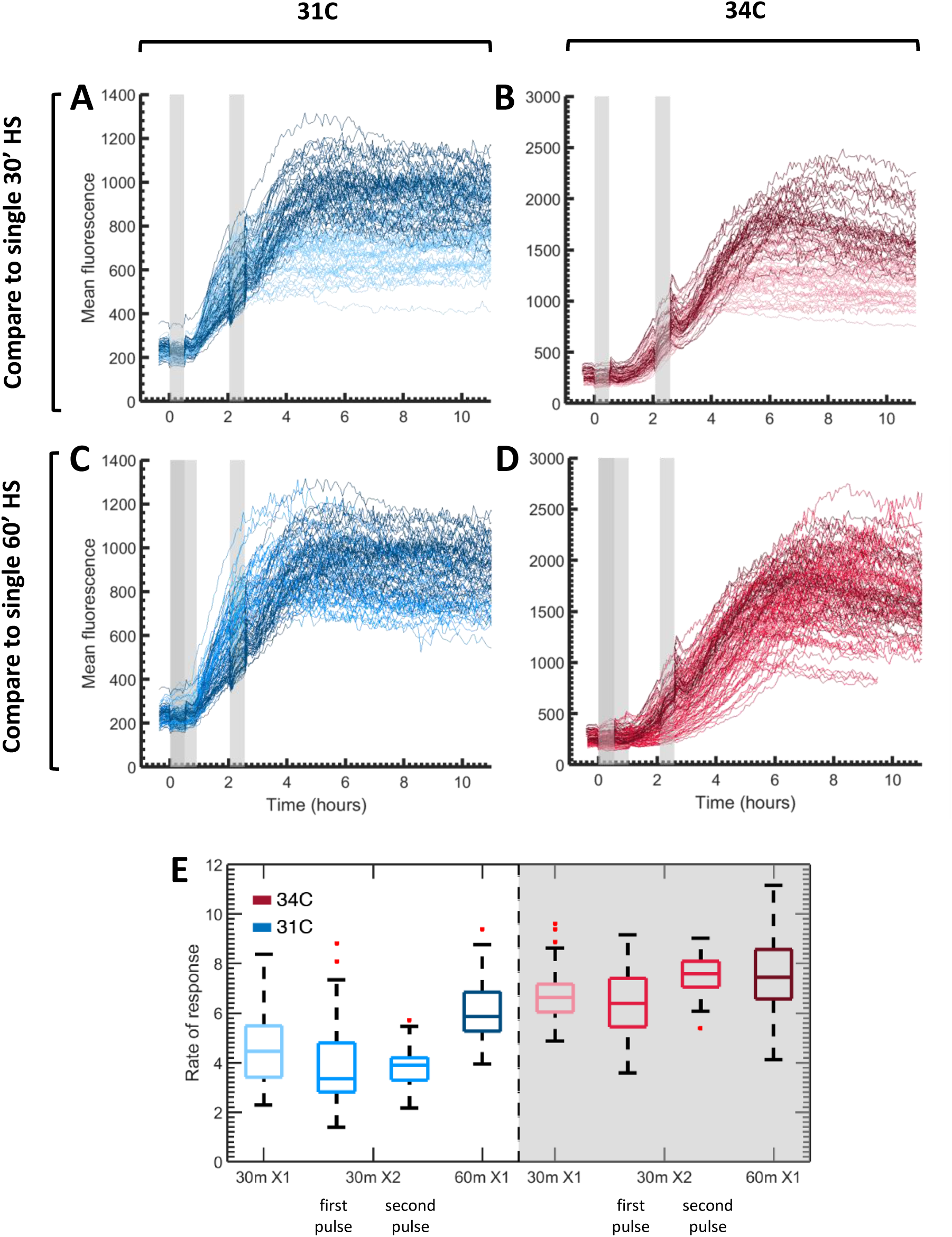
Extended data for two heat shock pulse experiments with wild type worms. **A, B** Individual response curves for worms subjected to a single 30 minute heat shock pulse (light color) or two consecutive 30 minute heat shock pulses separated by a 90 minute rest (dark color) at **(A)** 31°C (blue) or **(B)** 34°C (red). Heat shock pulse(s) indicated by grey bar(s). **C, D** Individual response curves for worms subjected to a single 60 minute heat shock pulse (medium color) or two consecutive 30 minute heat shock pulses separated by a 90 minute rest (dark color) at **(C)** 31°C (blue) or **(D)** 34°C (red). Heat shock pulse(s) indicated by grey bar(s). **E** Rates of response (mean fluorescence/minutes) for each pulse of two consecutive 30 minute heat shock pulses (medium color) compared to the rate of response for one single 30 minute (light color) or 60 minute (dark color) pulse at 31°C (blue) or 34°C (red).

**EV Figure 3.**
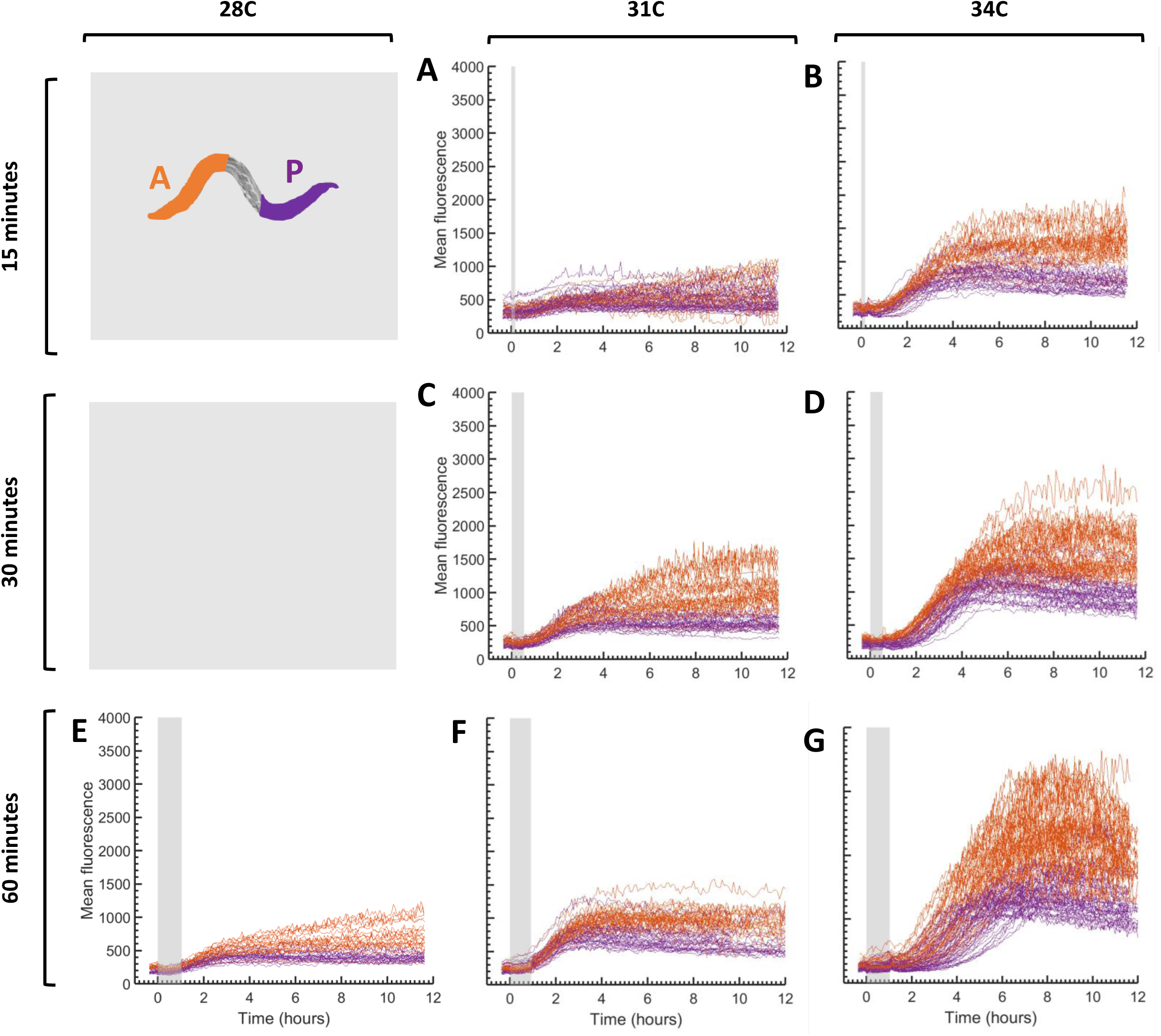
Individual worm anterior and posterior curves for single heat shock pulse experiments with wild type worms. **A – G** Individual response curves for anterior (orange) and posterior (purple) thirds of worm body for worms subject to a whole body heat shock pulse of **(A)** 15 minutes, 31°C; **(B)** 15 minutes, 34°C; **(C)** 30 minutes, 31°C; **(D)** 30 minutes, 34°C; **(E)** 60 minutes, 28°C; **(F)** 60 minutes, 31°C; and **(G)** 60 minutes, 34°C.

**EV Figure 4.**
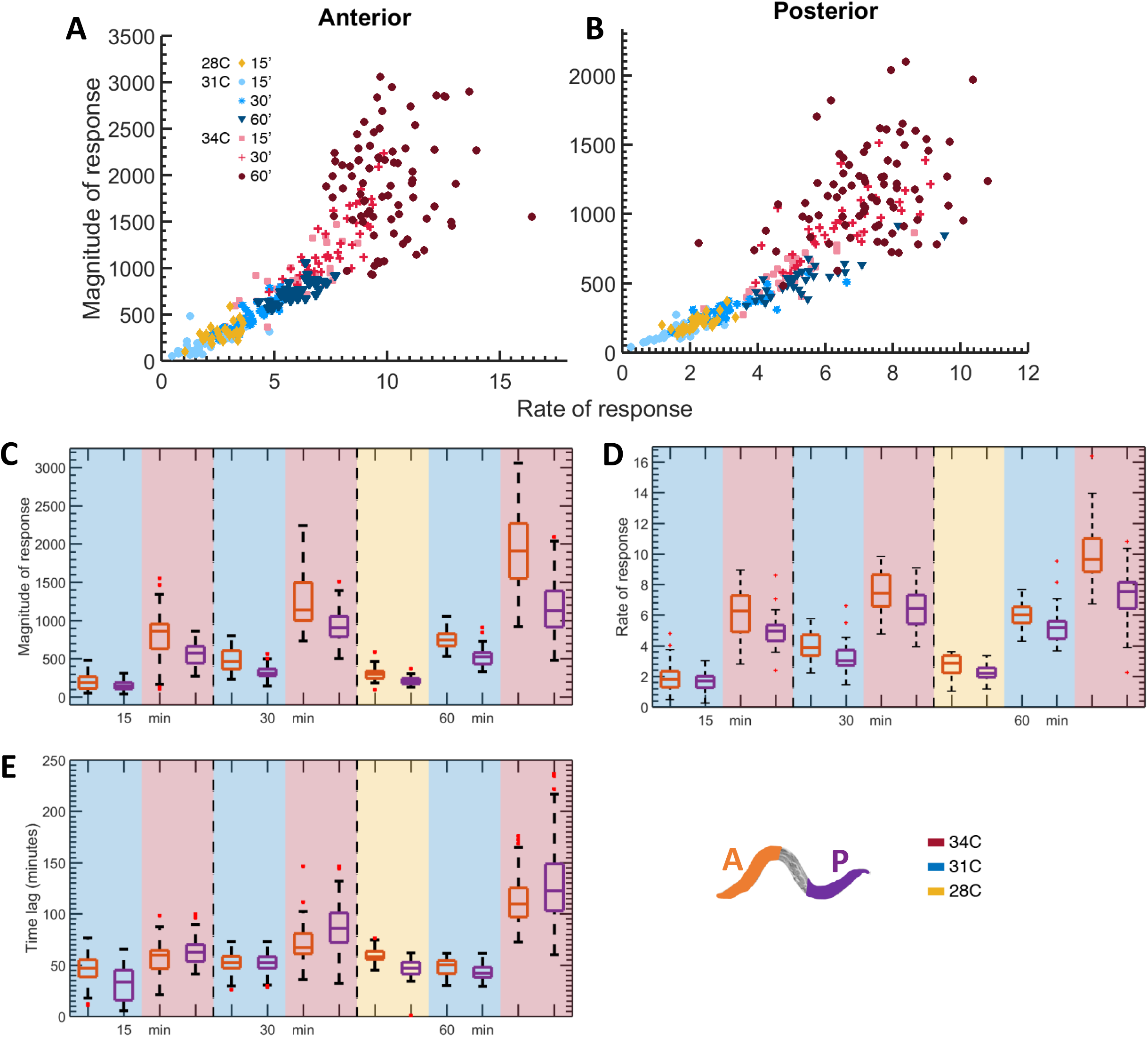
Extended data for spatial analysis of single heat shock pulse experiments with wild type worms. **A, B** Magnitude of response (mean fluorescence) plotted versus rate of response (mean fluorescence/minutes) for **(A)** anterior or **(B)** posterior part of worms subject to a whole-body, single heat shock pulse; duration and temperature as indicated in legend. **C, D (C)** Magnitude of response (mean fluorescence) and **(D)** rate of response (mean fluorescence/minutes) of anterior (orange) and posterior (purple) parts for 15, 30, or 60 minute heat shock pulse at 28°C (yellow), 31°C (blue), or 34°C (red). **E** Time lag (minutes) from start of heat shock pulse of anterior (orange) and posterior (purple) parts for 15, 30, or 60 minute heat shock pulse at 28°C (yellow), 31°C (blue), or 34°C (red).

**EV Figure 5.**
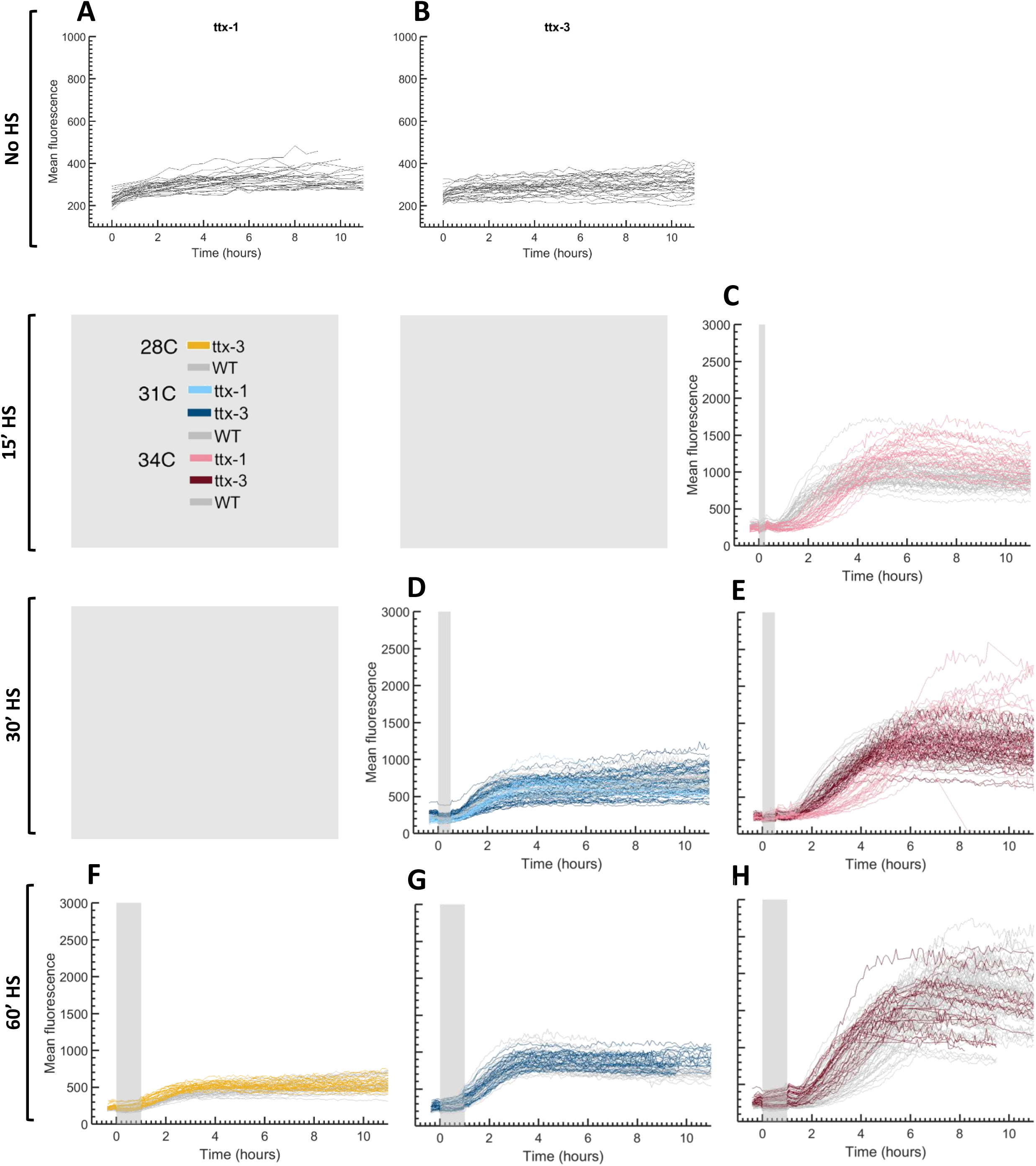
Individual worm curves for all single pulse heat shock experiments with *ttx-1* and *ttx-3* mutant worms. **A – H** Individual response curves for mutant worms subject to a whole body heat shock pulse of **(A)** no heat shock (*ttx-1*); **(B)** no heat shock (*ttx-3*); **(C)** 15 minutes, 34°C (*ttx-1*); **(D)** 30 minutes, 31°C (*ttx-1* & *ttx-3*); **(E)** 30 minutes, 34°C (*ttx-1* & *ttx-3*); **(F)** 60 minutes, 28°C (*ttx-3*); **(G)** 60 minutes, 31°C (*ttx-3*); and **(H)** 60 minutes, 34°C (*ttx-3*). Wild type worm curves for the same experimental conditions are indicated in grey for comparison. Each line indicates the mean fluorescence of an individual worm. Heat shock pulse is indicated by grey bar. Mean fluorescence was tracked for 10 hours following the heat shock pulse.

**EV Figure 6.**
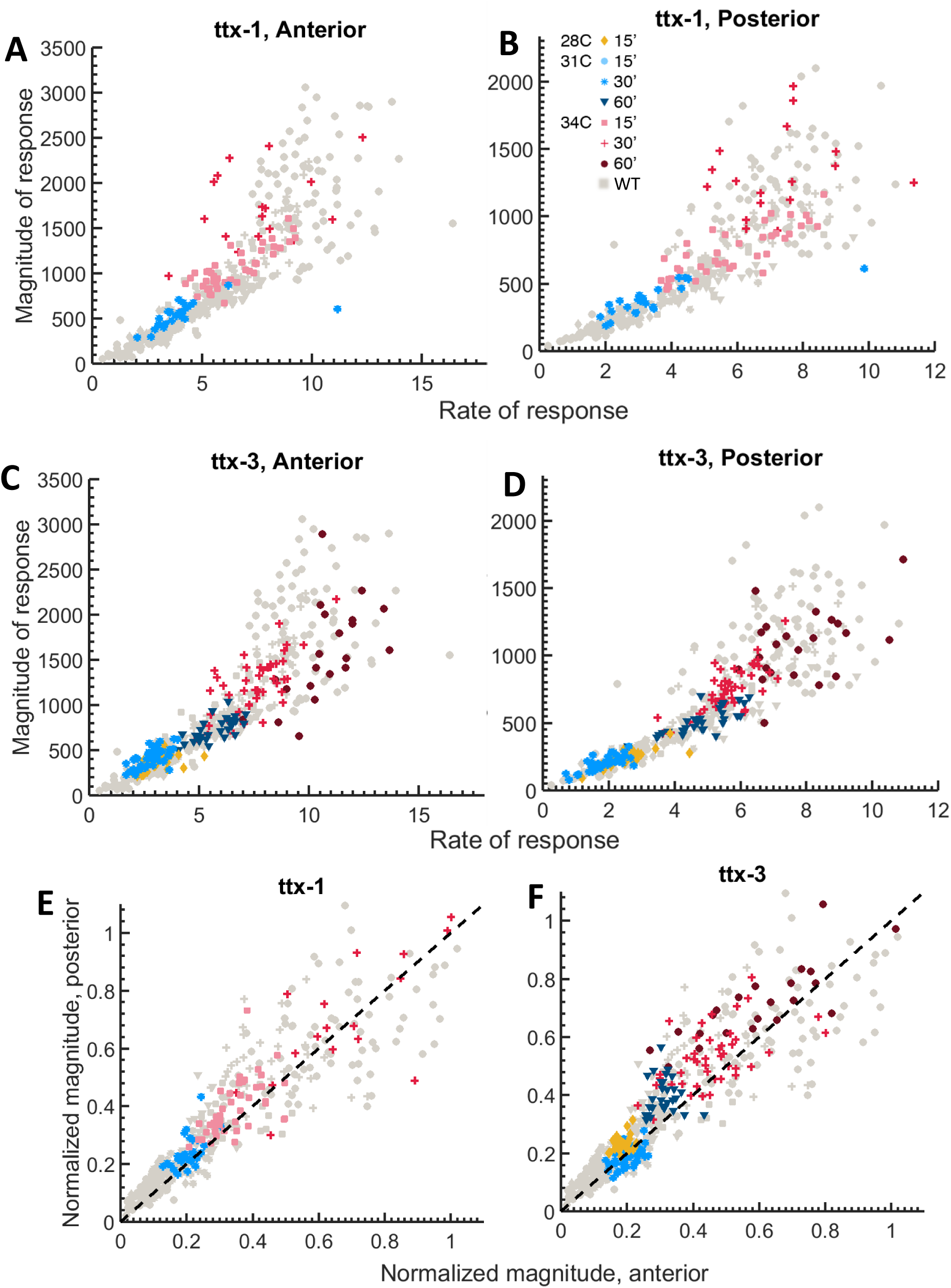
Extended data for *ttx-1* and *ttx-3* mutant single pulse heat shock experiments. **A, B** Magnitude of response (mean fluorescence) plotted versus rate of response (mean fluorescence/minutes) for **(A)** anterior or **(B)** posterior third of *ttx-1* (AFD) mutant worms subject to a whole-body, single heat shock pulse; duration and temperature are as indicated in legend. **C, D** Magnitude of response (mean fluorescence) plotted versus rate of response (mean fluorescence/minutes) for **(C)** anterior or **(D)** posterior third of *ttx-3* (AIY) mutant worms subject to a whole-body, single heat shock pulse; duration and temperature are as indicated in legend. **E, F** Magnitude of response (mean fluorescence) of posterior versus anterior of **(E)** *ttx-1* (AFD) and **(F)** *ttx-3* (AIY) mutant worms subject to a whole-body, single heat shock pulse; duration and temperature are as indicated in legend. Magnitudes are normalized to their respective maximum mean values in the anterior and posterior parts.

**EV Figure 7.**
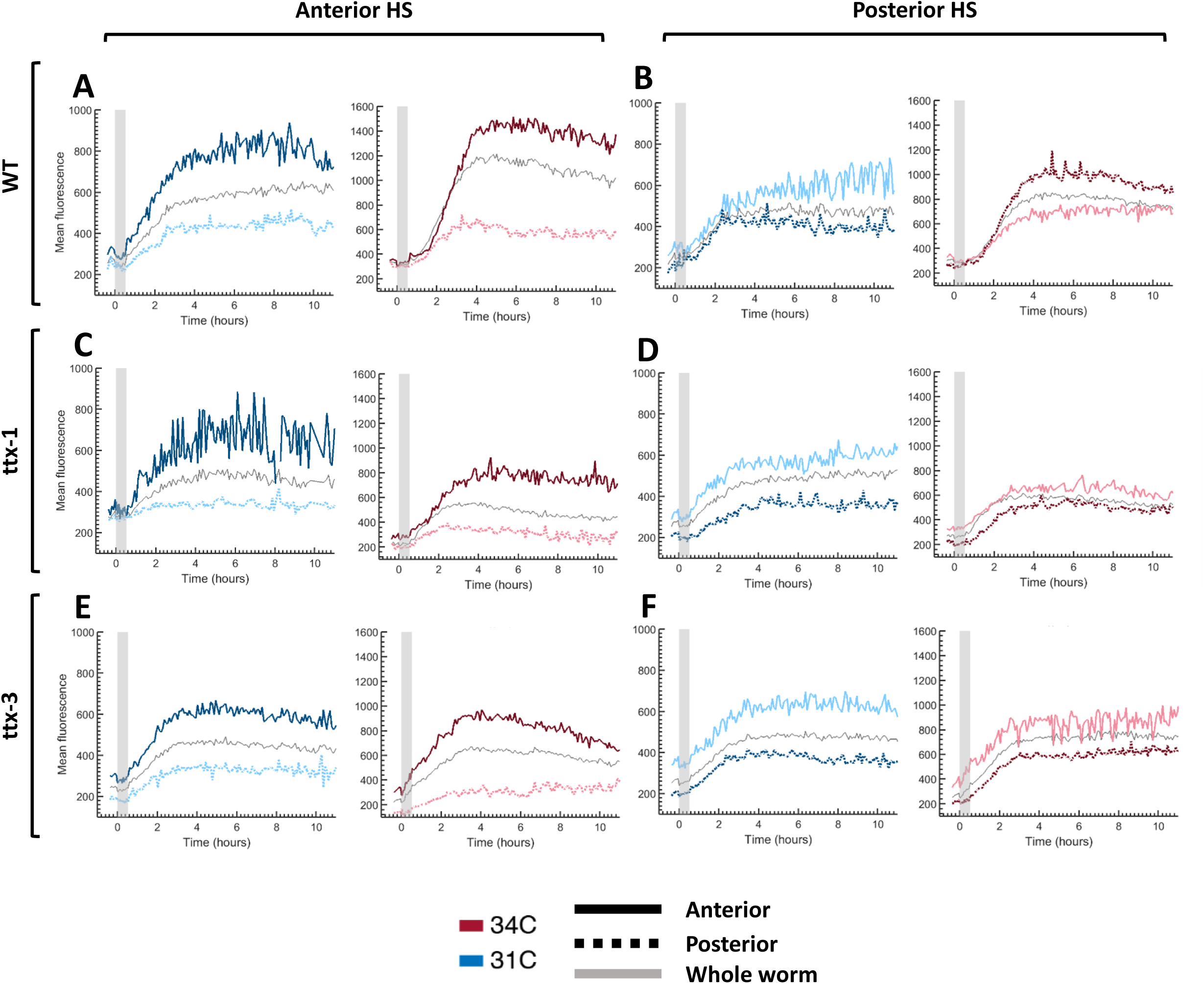
Example individual worm curves for 2 temperature heat shock experiments. **A – F** Example individual worm response curves for **(A, B)** wild type, **(C, D)** *ttx-1* (AFD) mutant worms, and **(E, F)** *ttx-3* (AIY) mutant worms subjected to a 30 minute **(A, C, E)** anterior-third-only or **(B, D, F)** posterior-third-only heat shock pulse at 31°C (blue) or 34°C (red). Grey line indicates mean fluorescence response averaged over the whole body. Darker line is mean fluorescence response averaged over the part of the body that was heat shocked. Lighter line is mean fluorescence response averaged over the part of the body that was kept at room temperature. Solid colored line is anterior; dashed colored line is posterior. Four columns going left to right are: (1) Heat shock anterior, 31°C; (2) Heat shock anterior, 34°C; (3) Heat shock posterior, 31°C; and (4) Heat shock posterior, 34°C.

**EV Figure 8.**
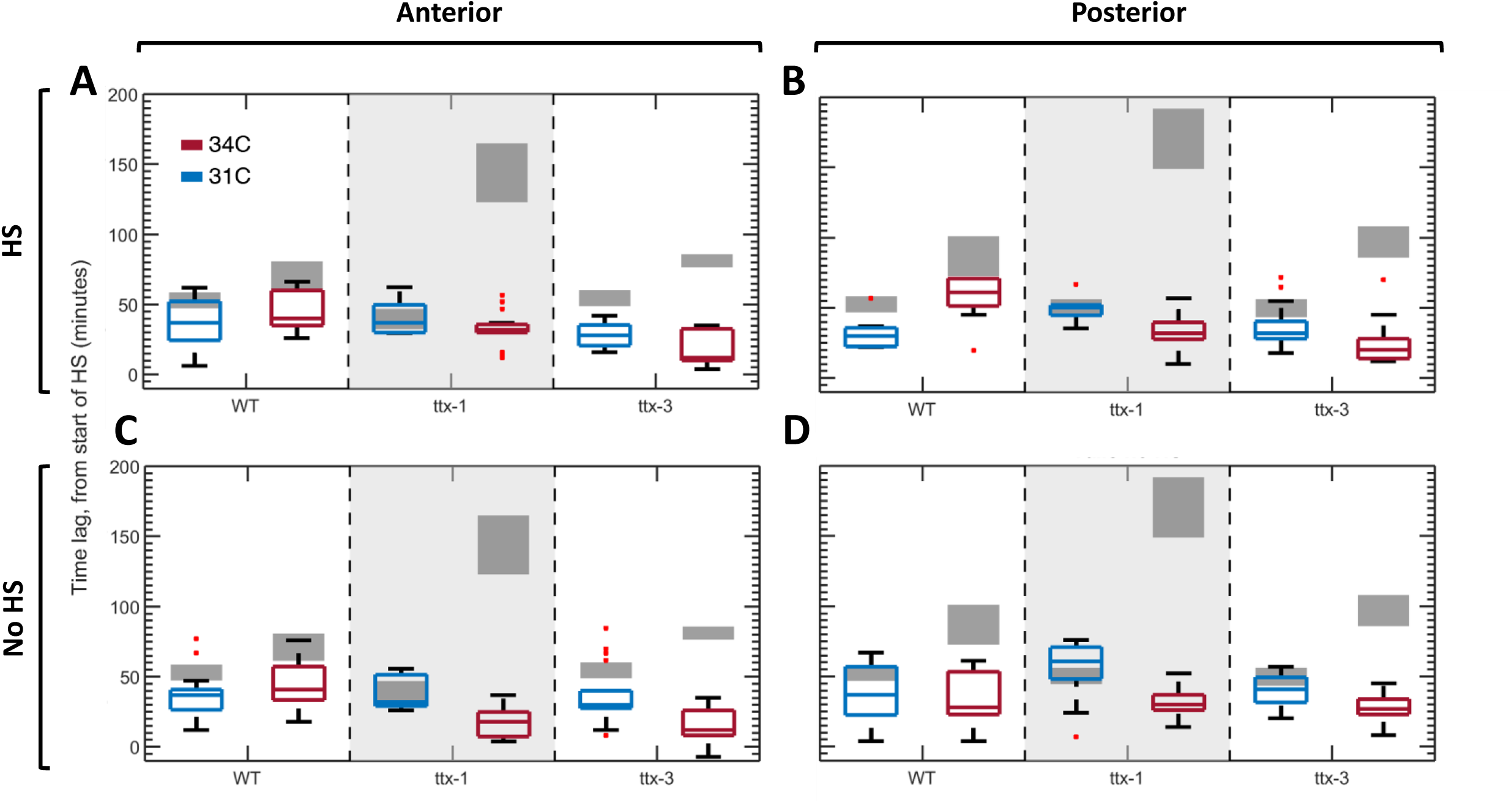
Time lag from start of heat shock for 2 temperature heat shock experiments. **A – D** Time lag in minutes from start of heat shock pulse for wild type, *ttx-1* (AFD) mutant, and *ttx-3* (AIY) mutant worms subjected to a 30 minute anterior- or posterior-third-only heat shock pulse at 31°C (blue) or 34°C (red). Lag is for **(A)** heat shock anterior, measure anterior; **(B)** heat shock posterior, measure posterior; **(C)** heat shock posterior, measure anterior; and **(D)** heat shock anterior, measure posterior. Grey boxes represent comparable anterior or posterior data for equivalent worm strain, given a full body heat shock pulse at the same temperature and duration.

**EV Video 1 | Example video of GFP fluorescence response of worm in microfluidics device.**

Example video of single wild type worm undergoing heat shock experiment in microfluidics device. Top video is phase contrast, middle video is GFP fluorescence, and bottom video is location of pixels in 95^th^ percentile of fluorescence and higher. Heat shock pulse is 60 minutes at 34°C and occurs from 0:20 – 1:20.

## Supporting Text

### Description of model and assumptions

We consider a simple mathematical model of the heat shock response (HSR). This model was inspired by previously-published models of the HSR in mammalian cells (Rieger et al., 2005; Scheff et al., 2015; Petre et al., 2011), but is considerably simplified to a minimal form that could still faithfully describe our experimental observations. Under normal conditions, the transcription factor (HSF) is located in a complex with a heat shock protein (HSP). Under stress, the complex dissociates, and the HSF is activated through a series of post-translational modifications, including trimerizing, translocating into the nucleus, and binding to a heat shock element on the DNA. For simplicity, we considered this activation as a single step. Activated HSF is used to induce transcription of the different hsp genes. In our model, we accounted for all of these genes collectively. In addition, HSF induces the gfp transgene in a similar manner. Our model assumes that the total concentration of HSF in all its forms does not change during the experiment. In contrast, all other moleculess undergo degradation. HSPs are also “used up” by forming a complex with a misfolded protein, produced during stress. Unused HSPs go back into complex with the HSF molecules, providing a negative feedback loop. This behavior is described by the following 8 equations:

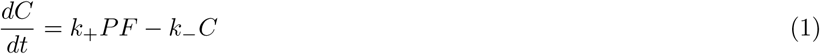

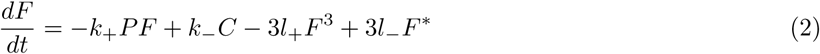

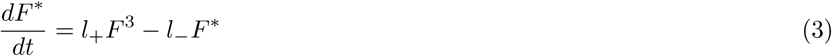

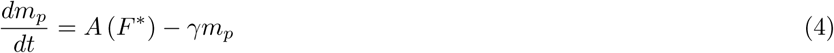

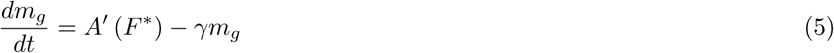

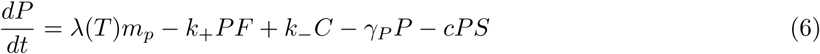

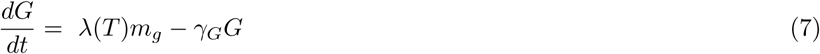

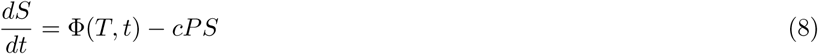

Here *C* is the concentration of the HSF:HSP complex; *F* and *F\** are the concentrations of free and active HSF respectively; *m_p_* and *m_g_* the concentrations of hsp and gfp mRNAs; *P* and *G* the concentrations of HSP and GFP, and *S* the concentration of misfolded proteins. The parameters are described in Table 1.

For simplicity, we assumed all mRNA have the same degradation rate. We also assumed they have the same transcription rate since they have the same promoter, and the same translation rate. However, we assumed significant basal transcription for HSP only. This reflects the basal amount of HSPs present under normal conditions, which does not include HSP-16.2. Activation of transcription by HSF is modeled in the form of a Hill function

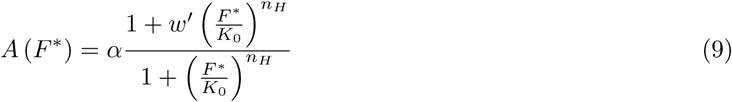

and *A*′ (*F\**) is modeled with the same form except for the 1 in the numerator.

Finally, we assumed that the interaction between HSPs and a misfolded protein is irreversible, mimicking their stoichiometric interaction. Accounting for partial recycling of HSPs only served to rescale some parameters of the model, with no significant effect on the results.

The behavior of the model can be described as follows. At steady state, a basal level of HSPs exists. Under low-level stress, this basal pool is not depleted, and the heat shock response will not be activated. This explains why the scale of stress level is a function of both the length and temperature of the heat shock, as both factors determine the consumption of HSPs. At intermediate stress levels, the basal level of HSPs is used up entirely, allowing a significant activation of HSF and induction of transcription of hsp mRNAs. Translation of the HSPs starts at the end of the heat shock, with the total translation rate depending on the accumulated concentration of mRNA. For a high stress dosage, the maximum number of mRNA possible will be produced, leading to the highest translation rate, and therefore the highest magnitude of response, measured by concentration of HSPs created. For a high enough stress, in order to deal with all of the misfolded proteins, the length of the response must then be increased. As the response ends, the HSP concentration will decrease back to its basal level.

### Quasi-steady state approximation

To simplify the model further, we first took advantage of the fact that the rates of HSF modifications (*<* seconds) are significantly faster than those of transcription/translation (minutes) and degradation (hours) by making quasi-steady state approximations for the HSF:HSP complex:

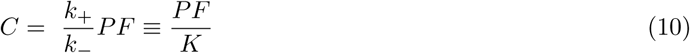

and for the activated form of HSF:

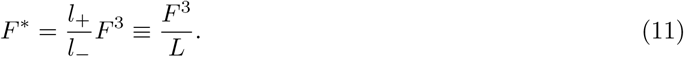

With these we rewrote the conservation law for the total HSF concentration,

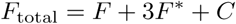

in the form

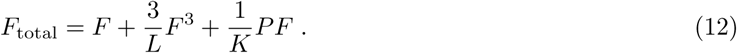

Relevant parameters can be found in Table 2. This equation relates the concentration of each form of HSF with the instantaneous concentration of HSPs.

### Final model

Combining these simplifications, we arrived at our final model:

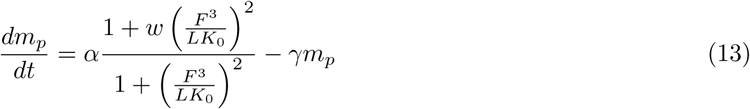

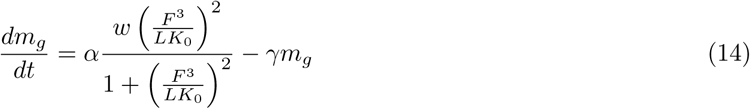

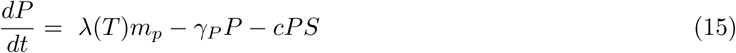

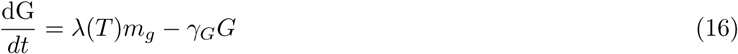

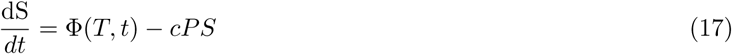

### Introducing delay in response to misfolded proteins

To explore how systemic response fits into this single cell model, we genetically ablated the AFD thermosen-sory neurons. We observed that this leads to a delay in the response and an increase in its magnitude, without affecting its rate. We therefore asked if a delay in the recognition of the stress signal could be the cause of the observed dynamics. To introduce such a time delay into the model, we added a delay to the stress signal that is recognized by HSPs, by replacing Equation (6) with

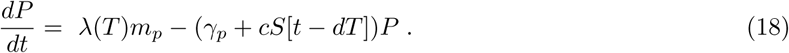

To match the observed data, *dT* was taken to be 1 hour.

### Tuning of parameter values

All values and units of parameters can be found in Table 3.

Degradation — mRNA in animals decays on a timescale of hours (Wang et al., 1999). Consistently, our qPCR results suggested a decay rate of 1 - 2 hours. GFP is known to be very stable in the worm intestine, and our data suggested a half-life of several days (Dietz and Rief, 2004). HSP degradation rate has very little effect on the dynamics of our model as long as it is fairly small, and even at larger values it only affected the steady state value of HSPs. We therefore took the half-life of both GFP and HSP to be on the order of 10s of hours.
Concentrations — We set the total HSF concentration (*F*_total_) to 1, and measured all other concentrations in comparison. In yeast and *Drosophila*, literature shows the steady state concentration of HSP is around 10 – 100 times larger than that of total HSF (Petre et al., 2011; Velazquez et al., 1983). Parameters were adjusted to set steady state ratios to this order of magnitude.
Transcription — Activation of HSR leads to induction of HSPs by 10 – 1000 fold (Wang et al., 1999; Sarge et al., 1993), and we chose *w* = 1000 to reflect the high end of this range. The order of magnitude of *K*_0_ was set by considering the activation dynamics we were looking for. Because of the form of the transcriptional term, *K*_0_*L* needed to be a little smaller than 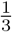 in order to be able to activate transcription. Similarly, *K* and *L* were defined relative to one another by considering the equilibrium state of HSF. To determine ranges of the translation and basal transcription rates, we considered the behavior we saw in our data. We wanted to saturate the rate of translation by saturating the number of mRNA made, not saturate the rate of transcription by saturating the number of active HSF.
Response — The rate constant *c*, which governs the kinetics of association between HSPs and misfolded protein, only affected the degree at which HSP production overshot its steady state level, with significant overshoot only for *c >* 1. Since our data indicated that such overshoot is only observed at high stress, we chose a marginal value of *c* = 1. The level of Φ at different temperatures determined the rate and magnitude of response. We picked these two values to best describe the measured kinetics at 31*^◦^* C and 34*^◦^* C. Once fixed, these values were kept for our entire analysis.

### Quantification of observed response dynamics

To quantify the observed expression dynamics of HSP-16.2p::GFP, we fit the observed fluorescence curves to the generalized logistic function

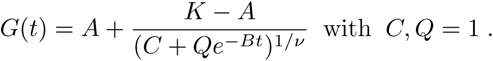

The fit was applied to a domain from the start of the experiment to around an hour after the maximum fluorescence was achieved. The end region of the experiment where rate of decline came into play was excluded for fitting purposes. We then used the fitted parameters to estimate the *Rate of response* as

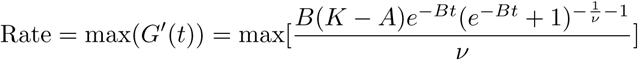

and the Magnitude of response as

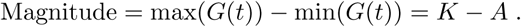

and the *Time lag* as

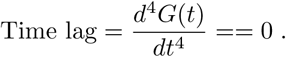

## Supporting Tables

**Supporting Table 1:** Model parameter definitions.

| Parameter | Definition |
| --- | --- |
| $k_+$ | Association rate of the HSF:HSP complex |
| $k_+$ | Dissociation rate of the HSF:HSP complex |
| $\ell_+$ | Activation rate of HSF |
| $\ell_-$ | Inactivation rate of HSF |
| $\gamma$ | mRNA degradation rate |
| $\gamma_P$ | Protein degradation rate of HSP |
| $\gamma_G$ | Protein degradation rate of GFP |
| $\lambda$ | Translation rate |
| $c$ | Association rate of HSP and a misfolded protein |
| $\Phi$ | Production rate of misfolded proteins |
| $w'$ | Maximal fold of activation |
| $K_0$ | Affinity of activated HSF to the hsp promoter |
| $\alpha$ | Basal transcription level |

**Supporting Table 2:** Parameters pertaining to Eq. 12.

| Parameter | Definition |
| --- | --- |
| $F_{\text{total}}$ | Total concentration of HSF |
| $L$ | Ratio of inactivation rate of HSF to activation rate |
| $K$ | Ratio of dissociation rate of the HSF:HSP complex to association rate |

**Supporting Table 3:** Estimated values of model parameters.

| Parameter | Value | Units |
| --- | --- | --- |
| $L$ | 0.1 | unitless |
| $K$ | 1 | unitless |
| $F_{\text{total}}$ | 1 | [F] |
| $\gamma$ | $\log(2)$ | hours<sup>-1</sup> |
| $\gamma_P$ | $\log(2)/100$ | hours<sup>-1</sup> |
| $\gamma_G$ | $\log(2)/50$ | hours<sup>-1</sup> |
| $\lambda(T)$ | 1 at RT, 0 otherwise | [P][m]<sup>-1</sup>hours<sup>-1</sup> |
| $c$ | 1 | [P]<sup>-1</sup>hours<sup>-1</sup> |
| $\Phi(T, t)$ | 2<sup>7</sup> at 31C, 2<sup>9</sup> at 34C | [S]hours<sup>-1</sup> |
| $w'$ | 1000 | unitless |
| $K_0$ | 0.1 | [F]<sup>3</sup> |
| $\alpha$ | 0.15 | [m]hours<sup>-1</sup> |

**Supporting Table 4:** Number of worms of each strain used in every experimental condition. Data for each condition is from at least 2 independent repeats.

| Experiment | WT | ttx-1 | ttx-3 |
| --- | --- | --- | --- |
| 15' @ 31°C | 38 | - | - |
| 15' @ 34°C | 51 | 30 | - |
| 30' @ 31°C | 42 | 28 | 42 |
| 30' @ 34°C | 41 | 28 | 48 |
| 60' @ 28°C | 34 | - | 32 |
| 60' @ 31°C | 37 | - | 29 |
| 60' @ 34°C | 83 | - | 30 |
| 2X 30' @ 31°C | 55 | - | - |
| 2X 30' @ 34°C | 38 | - | - |
| Anterior 30' @ 31°C | 19 | 21 | 12 |
| Posterior 30' @ 31°C | 15 | 6 | 30 |
| Anterior 30' @ 34°C | 16 | 19 | 13 |
| Posterior 30' @ 34°C | 21 | 13 | 24 |

## References

Abravaya K, Phillips B & Morimoto RI (1991) Heat shock-induced interactions of heat shock transcription factor and the human hsp70 promoter examined by in vivo footprinting. Mol. Cell. Biol. 11: 586–592

Ashburner M & Bonner JJ (1979) The induction of gene activity in drosophilia by heat shock. Cell 17: 241–254

Balch WE, Morimoto RI, Dillin A & Kelly JW (2008) Adapting Proteostasis for Disease Intervention. Science 319: 916–919

Balchin D, Hayer-Hartl M & Hartl FU (2016) In vivo aspects of protein folding and quality control. Science 353: aac4354

Bharadwaj S, Ali A & Ovsenek N (1999) Multiple Components of the HSP90 Chaperone Complex Function in Regulation of Heat Shock Factor 1 In Vivo. Mol. Cell. Biol. 19: 8033–8041

Bose S & Cho J (2016) Targeting chaperones, heat shock factor-1, and unfolded protein response: Promising therapeutic approaches for neurodegenerative disorders. Ageing Res. Rev.

Bouche G, Amalric F, Caizergues-Ferrer M & Zalta JP (1979) Effects of heat shock on gene expression and subcellular protein distribution in Chinese hamster ovary cells. Nucleic Acids Res. 7: 1739–1747

Castells-Roca L, García-Martínez J, Moreno J, Herrero E, Bellí G & Pérez-Ortín JE (2011) Heat Shock Response in Yeast Involves Changes in Both Transcription Rates and mRNA Stabilities. PLoS ONE 6: e17272

Cates J, Graham GC, Omattage N, Pavesich E, Setliff I, Shaw J, Smith C & Lipan O (2011) Sensing the Heat Stress by Mammalian Cells. BMC Biophys. 4: 16

Chiang W-C, Ching T-T, Lee HC, Mousigian C & Hsu A-L (2012) HSF-1 Regulators DDL-1/2 Link Insulin-like Signaling to Heat-Shock Responses and Modulation of Longevity. Cell 148: 322–334

Dietz H & Rief M (2004) Exploring the energy landscape of GFP by single-molecule mechanical experiments. Proc. Natl. Acad. Sci. U. S. A. 101: 16192–16197

Durieux J, Wolff S & Dillin A (2011) The Cell-Non-Autonomous Nature of Electron Transport Chain-Mediated Longevity. Cell 144: 79–91

El-Samad H, Kurata H, Doyle JC, Gross CA & Khammash M (2005) Surviving heat shock: Control strategies for robustness and performance. Proc. Natl. Acad. Sci. U. S. A. 102: 2736–2741

Feder ME & Hofmann GE (1999) HEAT-SHOCK PROTEINS, MOLECULAR CHAPERONES, AND THE STRESS RESPONSE: Evolutionary and Ecological Physiology. Annu. Rev. Physiol. 61: 243–282

Finka A, Sood V, Quadroni M, Rios PDL & Goloubinoff P (2015) Quantitative proteomics of heat-treated human cells show an across-the-board mild depletion of housekeeping proteins to massively accumulate few HSPs. Cell Stress Chaperones 20: 605–620

Fritsch M & Wu C (1999) Phosphorylation of Drosophila heat shock transcription factor. Cell Stress Chaperones 4: 102–117

Fukuoka M, Yoshida M, Eda A, Takahashi M & Hohjoh H (2014) Gene Silencing Mediated by Endogenous MicroRNAs under Heat Stress Conditions in Mammalian Cells. PloS One 9: e103130

Garcia SM, Casanueva MO, Silva MC, Amaral MD & Morimoto RI (2007) Neuronal signaling modulates protein homeostasis in Caenorhabditis elegans post-synaptic muscle cells. Genes Dev. 21: 3006–3016

Garigan D, Hsu A-L, Fraser AG, Kamath RS, Ahringer J & Kenyon C (2002) Genetic Analysis of Tissue Aging in Caenorhabditis elegans: A Role for Heat-Shock Factor and Bacterial Proliferation. Genetics 161: 1101– 1112

Gidalevitz T, Prahlad V & Morimoto RI (2011) The Stress of Protein Misfolding: From Single Cells to Multicellular Organisms. Cold Spring Harb. Perspect. Biol. 3: a009704–a009704

Gruber J, Tang SY & Halliwell B (2007) Evidence for a trade-off between survival and fitness caused by resveratrol treatment of Caenorhabditis elegans. Ann. N. Y. Acad. Sci. 1100: 530–542

Guisbert E, Czyz DM, Richter K, McMullen PD & Morimoto RI (2013) Identification of a Tissue-Selective Heat Shock Response Regulatory Network. PLoS Genet 9: e1003466

Guisbert E, Yura T, Rhodius VA & Gross CA (2008) Convergence of Molecular, Modeling, and Systems Approaches for an Understanding of the Escherichia coli Heat Shock Response. Microbiol. Mol. Biol. Rev. 72: 545–554

Guo Y, Guettouche T, Fenna M, Boellmann F, Pratt WB, Toft DO, Smith DF & Voellmy R (2001) Evidence for a Mechanism of Repression of Heat Shock Factor 1 Transcriptional Activity by a Multichaperone Complex. J. Biol. Chem. 276: 45791–45799

Hobert O, Mori I, Yamashita Y, Honda H, Ohshima Y, Liu Y & Ruvkun G (1997) Regulation of Interneuron Function in the C. elegans Thermoregulatory Pathway by the ttx-3 LIM Homeobox Gene. Neuron 19: 345– 357

Inoue M, Mitarai N & Trusina A (2012) Circuit architecture explains functional similarity of bacterial heat shock responses. Phys. Biol. 9: 066003

Kang H-W (2012) A multiscale approximation in a heat shock response model of E. coli. BMC Syst. Biol. 6: 143

Kantidze OL, Velichko AK, Luzhin AV & Razin SV (2016) Heat Stress-Induced DNA Damage. Acta Naturae 8: 75–78

Kline MP & Morimoto RI (1997) Repression of the heat shock factor 1 transcriptional activation domain is modulated by constitutive phosphorylation. Mol. Cell. Biol. 17: 2107–2115

Kopito RB & Levine E (2014) Durable spatiotemporal surveillance of Caenorhabditis elegans response to environmental cues. Lab. Chip 14: 764–770

Krakowiak J, Zheng X, Patel N, Anandhakumar J, Valerius K, Gross DS, Khalil AS & Pincus D (2017) Hsf1 and Hsp70 constitute a two-component feedback loop that regulates the yeast heat shock response. bioRxiv: 183137

Kumsta C, Ching T-T, Nishimura M, Davis AE, Gelino S, Catan HH, Yu X, Chu C-C, Ong B, Panowski SH, Baird N, Bodmer R, Hsu A-L & Hansen M (2014) Integrin-linked kinase modulates longevity and thermotolerance in C. elegans through neuronal control of HSF-1. Aging Cell 13: 419–430

Kurata H, El-Samad H, Iwasaki R, Ohtake H, Doyle JC, Grigorova I, Gross CA & Khammash M (2006) Module-Based Analysis of Robustness Tradeoffs in the Heat Shock Response System. PLoS Comput Biol 2: e59

Leach MD, Tyc KM, Brown AJP & Klipp E (2012) Modelling the Regulation of Thermal Adaptation in Candida albicans, a Major Fungal Pathogen of Humans. PLoS ONE 7: e32467

Lee S-J & Kenyon C (2009) Regulation of the Longevity Response to Temperature by Thermosensory Neurons in Caenorhabditis elegans. Curr. Biol. 19: 715–722

Leiser SF, Rossner R & Kaeberlein M (2016) New insights into cell non-autonomous mechanisms of the C. elegans hypoxic response. Worm 5: e1176823

Lindquist S (1980) Varying patterns of protein synthesis in Drosophila during heat shock: Implications for regulation. Dev. Biol. 77: 463–479

Link CD, Cypser JR, Johnson CJ & Johnson TE (1999) Direct Observation of Stress Response in Caenorhabditis elegans Using a Reporter Transgene. Cell Stress Chaperones 4: 235–242

Luo L, Cook N, Venkatachalam V, Martinez-Velazquez LA, Zhang X, Calvo AC, Hawk J, MacInnis BL, Frank M, Ng JHR, Klein M, Gershow M, Hammarlund M, Goodman MB, Colón-Ramos DA, Zhang Y & Samuel ADT (2014) Bidirectional thermotaxis in Caenorhabditis elegans is mediated by distinct sensorimotor strategies driven by the AFD thermosensory neurons. Proc. Natl. Acad. Sci. U. S. A. 111: 2776–2781

Ma J, Grant CE, Plagens RN, Barrett LN, Kim Guisbert KS & Guisbert E (2017) Cellular Proteomes Drive Tissue-Specific Regulation of the Heat Shock Response. G3 Bethesda Md

Maxwell CS, Kruesi WS, Core LJ, Kurhanewicz N, Waters CT, Lewarch CL, Antoshechkin I, Lis JT, Meyer BJ & Baugh LR (2014) Pol II Docking and Pausing at Growth and Stress Genes in C. elegans. Cell Rep. 6: 455– 466

McCallum KC, Liu B, Fierro-González JC, Swoboda P, Arur S, Miranda-Vizuete A & Garsin DA (2016) TRX-1 Regulates SKN-1 Nuclear Localization Cell Non-autonomously in Caenorhabditis elegans. Genetics

Mendenhall AR, Tedesco PM, Sands B, Johnson TE & Brent R (2015) Single Cell Quantification of Reporter Gene Expression in Live Adult Caenorhabditis elegans Reveals Reproducible Cell-Specific Expression Patterns and Underlying Biological Variation. PloS One 10: e0124289

Mendenhall AR, Tedesco PM, Taylor LD, Lowe A, Cypser JR & Johnson TE (2012) Expression of a Single-Copy hsp-16.2 Reporter Predicts Life span. J. Gerontol. A. Biol. Sci. Med. Sci. Available at: http://biomedgerontology.oxfordjournals.org/content/early/2012/01/06/gerona.glr225 [Accessed May 1, 2013]

Mendillo ML, Santagata S, Koeva M, Bell GW, Hu R, Tamimi RM, Fraenkel E, Ince TA, Whitesell L & Lindquist S (2012) HSF1 drives a transcriptional program distinct from heat shock to support highly malignant human cancers. Cell 150: 549–562

Mori I & Ohshima Y (1995a) Neural regulation of thermotaxis in Caenorhabditis elegans. Nature 376: 344–348

Mori I & Ohshima Y (1995b) Neural regulation of thermotaxis in Caenorhabditis elegans. Nature 376: 344–348

Morimoto RI (2008) Proteotoxic stress and inducible chaperone networks in neurodegenerative disease and aging. Genes Dev. 22: 1427–1438

Morimoto RI (2011) The Heat Shock Response: Systems Biology of Proteotoxic Stress in Aging and Disease. Cold Spring Harb. Symp. Quant. Biol. 76: 91–99

Niskanen EA, Malinen M, Sutinen P, Toropainen S, Paakinaho V, Vihervaara A, Joutsen J, Kaikkonen MU, Sistonen L & Palvimo JJ (2015) Global SUMOylation on active chromatin is an acute heat stress response restricting transcription. Genome Biol. 16: 153

Nussbaum-Krammer CI & Morimoto RI (2014) Caenorhabditis elegans as a model system for studying non-cell-autonomous mechanisms in protein-misfolding diseases. Dis. Model. Mech. 7: 31–39

Ohama N, Sato H, Shinozaki K & Yamaguchi-Shinozaki K (2016) Transcriptional Regulatory Network of Plant Heat Stress Response. Trends Plant Sci.

van Oosten-Hawle P, Porter RS & Morimoto RI (2013) Regulation of organismal proteostasis by transcellular chaperone signaling. Cell 153: 1366–1378

Petre I, Mizera A, Hyder CL, Meinander A, Mikhailov A, Morimoto RI, Sistonen L, Eriksson JE & Back R-J (2011) A simple mass-action model for the eukaryotic heat shock response and its mathematical validation. Nat. Comput. 10: 595–612

Powers ET, Morimoto RI, Dillin A, Kelly JW & Balch WE (2009) Biological and Chemical Approaches to Diseases of Proteostasis Deficiency. Annu. Rev. Biochem. 78: 959–991

Prahlad V, Cornelius T & Morimoto RI (2008) Regulation of the Cellular Heat Shock Response in Caenorhabditis elegans by Thermosensory Neurons. Science 320: 811–814

Prahlad V & Morimoto RI (2009) Integrating the stress response: lessons for neurodegenerative diseases from C. elegans. Trends Cell Biol. 19: 52–61

Raynes R, Leckey BD, Nguyen K & Westerheide SD (2012) Heat Shock and Caloric Restriction Have a Synergistic Effect on the Heat Shock Response in a sir2.1-dependent Manner in Caenorhabditis elegans. J. Biol. Chem. 287: 29045–29053

Rieger TR, Morimoto RI & Hatzimanikatis V (2005) Mathematical Modeling of the Eukaryotic Heat-Shock Response: Dynamics of the hsp70 Promoter. Biophys. J. 88: 1646–1658

Rieger TR, Morimoto RI & Hatzimanikatis V (2006) Bistability Explains Threshold Phenomena in Protein Aggregation both In Vitro and In Vivo. Biophys. J. 90: 886–895

Sarge KD, Murphy SP & Morimoto RI (1993) Activation of heat shock gene transcription by heat shock factor 1 involves oligomerization, acquisition of DNA-binding activity, and nuclear localization and can occur in the absence of stress. Mol. Cell. Biol. 13: 1392–1407

Satterlee JS, Sasakura H, Kuhara A, Berkeley M, Mori I & Sengupta P (2001) Specification of Thermosensory Neuron Fate in C. elegans Requires ttx-1, a Homolog of otd/Otx. Neuron 31: 943–956

Satyal SH, Schmidt E, Kitagawa K, Sondheimer N, Lindquist S, Kramer JM & Morimoto RI (2000) Polyglutamine aggregates alter protein folding homeostasis in Caenorhabditis elegans. Proc. Natl. Acad. Sci. 97: 5750– 5755

Scheff JD, Stallings JD, Reifman J & Rakesh V (2015) Mathematical Modeling of the Heat-Shock Response in HeLa Cells. Biophys. J. 109: 182–193

Shalgi R, Hurt JA, Krykbaeva I, Taipale M, Lindquist S & Burge CB (2013) Widespread Regulation of Translation by Elongation Pausing in Heat Shock. Mol. Cell 49: 439–452

Shi Y, Mosser DD & Morimoto RI (1998) Molecularchaperones as HSF1-specific transcriptional repressors. Genes Dev. 12: 654–666

Sivéry A, Courtade E & Thommen Q (2016) A minimal titration model of the mammalian dynamical heat shock response. Phys. Biol. 13: 066008

Somero GN (1995) Proteins and Temperature. Annu. Rev. Physiol. 57: 43–68

Soukup ST, Spanier B, Grünz G, Bunzel D, Daniel H & Kulling SE (2012) Formation of phosphoglycosides in Caenorhabditis elegans: a novel biotransformation pathway. PloS One 7: e46914

Spriggs KA, Bushell M & Willis AE (2010) Translational Regulation of Gene Expression during Conditions of Cell Stress. Mol. Cell 40: 228–237

Sriram K, Rodriguez-Fernandez M & Iii FJD (2012) A Detailed Modular Analysis of Heat-Shock Protein Dynamics under Acute and Chronic Stress and Its Implication in Anxiety Disorders. PLOS ONE 7: e42958

Starosta AL, Lassak J, Jung K & Wilson DN (2014) The bacterial translation stress response. FEMS Microbiol. Rev.

Stiernagle T (2006) Maintenance of C. elegans. WormBook Available at: http://www.wormbook.org/chapters/www_strainmaintain/strainmaintain.html [Accessed July 7, 2013]

Su K-H, Cao J, Tang Z, Dai S, He Y, Sampson SB, Benjamin IJ & Dai C (2016) HSF1 critically attunes proteotoxic stress sensing by mTORC1 to combat stress and promote growth. Nat. Cell Biol.

Takeuchi T, Suzuki M, Fujikake N, Popiel HA, Kikuchi H, Futaki S, Wada K & Nagai Y (2015) Intercellular chaperone transmission via exosomes contributes to maintenance of protein homeostasis at the organismal level. Proc. Natl. Acad. Sci. U. S. A. 112: E2497–2506

Tatum MC, Ooi FK, Chikka MR, Chauve L, Martinez-Velazquez LA, Steinbusch HWM, Morimoto RI & Prahlad V (2015) Neuronal Serotonin Release Triggers the Heat Shock Response in C. elegans in the Absence of Temperature Increase. Curr. Biol. 25: 163–174

Taylor RC & Dillin A (2013) XBP-1 Is a Cell-Nonautonomous Regulator of Stress Resistance and Longevity. Cell 153: 1435–1447

Vabulas RM, Raychaudhuri S, Hayer-Hartl M & Hartl FU (2010) Protein Folding in the Cytoplasm and the Heat Shock Response. Cold Spring Harb. Perspect. Biol. 2: a004390

Velazquez JM, Sonoda S, Bugaisky G & Lindquist S (1983) Is the major Drosophila heat shock protein present in cells that have not been heat shocked? J. Cell Biol. 96: 286–290

Wang SM, Khandekar JD, Kaul KL, Winchester DJ & Morimoto RI (1999) A method for the quantitative analysis of human heat shock gene expression using a multiplex RT-PCR assay. Cell Stress Chaperones 4: 153–161

Westerheide SD & Morimoto RI (2005) Heat shock response modulators as therapeutic tools for diseases of protein conformation. J. Biol. Chem. 280: 33097–33100

Wolff S, Weissman JS & Dillin A (2014) Differential Scales of Protein Quality Control. Cell 157: 52–64

Zander G, Hackmann A, Bender L, Becker D, Lingner T, Salinas G & Krebber H (2016) mRNA quality control is bypassed for immediate export of stress-responsive transcripts. Nature

Zou J, Guo Y, Guettouche T, Smith DF & Voellmy R (1998) Repression of Heat Shock Transcription Factor HSF1 Activation by HSP90 (HSP90 Complex) that Forms a Stress-Sensitive Complex with HSF1. Cell 94: 471– 480

Zullo J, Matsumoto K, Xavier S, Ratliff B & Goligorsky MS (2015) The cell secretome, a mediator of cell-to-cell communication. Prostaglandins Other Lipid Mediat.

## References

Hendrik Dietz and Matthias Rief. Exploring the energy landscape of GFP by single-molecule mechanical experiments. Proceedings of the National Academy of Sciences of the United States of America, 101(46): 16192–16197, November 2004.

Ion Petre, Andrzej Mizera, Claire L. Hyder, Annika Meinander, Andrey Mikhailov, Richard I. Morimoto, Lea Sistonen, John E. Eriksson, and Ralph-Johan Back. A simple mass-action model for the eukaryotic heat shock response and its mathematical validation. Natural Computing, 10(1):595–612, March 2011.

Theodore R. Rieger, Richard I. Morimoto, and Vassily Hatzimanikatis. Mathematical modeling of the eukaryotic heat-shock response: dynamics of the hsp70 promoter. Biophysical Journal, 88(3):1646–1658, March 2005.

K D Sarge, S P Murphy, and R I Morimoto. Activation of heat shock gene transcription by heat shock factor 1 involves oligomerization, acquisition of DNA-binding activity, and nuclear localization and can occur in the absence of stress. Molecular and Cellular Biology, 13(3):1392–1407, March 1993.

Jeremy D. Scheff, Jonathan D. Stallings, Jaques Reifman, and Vineet Rakesh. Mathematical Modeling of the Heat-Shock Response in HeLa Cells. Biophysical Journal, 109(2):182–193, July 2015.

J. M. Velazquez, S. Sonoda, G. Bugaisky, and S. Lindquist. Is the major Drosophila heat shock protein present in cells that have not been heat shocked? The Journal of Cell Biology, 96(1):286–290, January 1983.

San Ming Wang, Janardan D. Khandekar, Karen L. Kaul, David J. Winchester, and Richard I. Morimoto. A method for the quantitative analysis of human heat shock gene expression using a multiplex RT-PCR assay. Cell Stress & Chaperones, 4(3):153–161, September 1999.

